# Structural basis for a natural circular permutation in proteins

**DOI:** 10.1101/2020.10.28.360099

**Authors:** Samuel G. Nonis, Joel Haywood, Jason W. Schmidberger, Charles S. Bond, Joshua S. Mylne

**Affiliations:** The University of Western Australia, School of Molecular Sciences, 35 Stirling Highway, Crawley, Perth 6009, Australia; The University of Western Australia, The ARC Centre of Excellence in Plant Energy Biology, 35 Stirling Highway, Crawley, Perth 6009, Australia

## Abstract

Over 30 years ago, an intriguing post-translational modification was discovered to be responsible for creating concanavalin A (conA), a carbohydrate-binding protein found in the seeds of jack bean (*Canavalia ensiformis*) and commercially used for carbohydrate chromatography. Biosynthesis of conA involves what was then an unprecedented rearrangement in amino acid sequence, whereby the N-terminal half of the gene-encoded conA precursor is swapped to become the C-terminal half of conA. The cysteine protease, asparaginyl endopeptidase (AEP), was shown to be involved, but its mechanism was not fully elucidated. To understand the structural basis and consequences of conA circular permutation, we generated a recombinant jack bean conA precursor (pro-conA) plus jack bean AEP (CeAEP1) and solved crystal structures for each to 2.1 Å and 2.7 Å respectively. By reconstituting the biosynthesis of conA *in vitro*, we prove CeAEP1 alone can perform both the cleavage and cleavage-coupled transpeptidation to form conA. CeAEP1 structural analysis reveals how it is capable of carrying out both these reactions. Biophysical assays illustrated that conA is more thermally and pH stable than pro-conA, consistent with fewer intermolecular interactions between subunits in the pro-conA crystal structure. These findings elucidate the consequences of circular permutation in the only post-translation example known to occur in nature.

## Introduction

Concanavalin A (conA) is a seed lectin of the jack bean plant (*Canavalia ensiformis*); it is a non-catalytic protein that binds specific carbohydrates (monomers and oligomers of mannose and glucose) reversibly and with high specificity and moderate affinity (Lis and Sharon, 1998). Since its discovery just over a century ago (Sumner, 1919), conA has become the most studied lectin largely due to interest in its carbohydrate-binding properties *in vitro* (Bernhard and Avrameas, 1971; Dwyer and Johnson, 1981; Goldstein et al., 1997; Lis and Sharon, 1998; Locke et al., 2014), and also due to the unusual post-translational circular permutation it undergoes to become the mature form. The carbohydrate binding of conA has seen it widely adopted in chromatography where it is frequently immobilised on sepharose and used to purify glycosylated biomolecules bearing high-mannose type N-glycans, including glycoproteins, polysaccharides and glycolipids (Ogata et al., 1975; Saleemuddin and Husain, 1991). Despite a depth of structural knowledge, evidenced by over 60 jack bean conA structures in complex with various ligands and metal ions which are necessary for its function, the properties of pro-conA and its maturation into conA are not fully understood.

The hypothesised biological roles of conA include involvement in seed storage and plant defence. These are based mostly on its ability to bind certain carbohydrates, and are often inferred from conclusions drawn from studies performed on other legume lectins with similar physico-chemical properties (Sharon and Lis, 1990). Lectins have been suggested to function as packaging aids as they are associated with other storage proteins in the developing protein bodies of seeds (Einhoff et al., 1986). A lowering of pH during water imbibition was postulated to help lectins dissociate from storage proteins, possibly allowing lectins to diffuse out of seedlings and contribute to protection against bacterial, fungal, and viral pathogens (Peumans and Van Damme, 1995). A major argument for the role of lectins in plant immunity is based on their interactions with glyco-components that are absent in plants but are found on the surface of microbes or along the digestive tract of insects and animals (Peumans and Van Damme, 1995; Lagarda-Diaz et al., 2017). Several lectins have indeed been shown to be resistant to proteolysis by digestive enzymes and to have insecticidal properties (Melander et al., 2003; Macedo et al., 2007). As conA is synthesised with an asparagine-linked (N-linked) glycan that inhibits its carbohydrate-binding ability, but can interact with jack bean storage proteins and plant growth hormones in the mature form *in vitro*, it seems likely that conA function is spatially and/or temporally regulated (Edelman and Wang, 1978; Smith et al., 1982). With conA making up about 20% of the jack bean seed storage protein content {Dalkin, 1983 #10343}, it could function not only as a potent passive defensive mechanism in the metabolically inactive seed, but also as a source of amino acids and metal ions during germination.

The biosynthesis of conA involves a unique series of cleavages and a transpeptidation reaction that occur on the carboxyl side of asparagine residues, resulting in conA circular permutation (**Figure 1A**). ConA is synthesised as an inactive glycoprotein precursor (pre-pro-conA) (Herman et al., 1985) (**Supplemental Figure 1**). The N-linked glycan inhibits carbohydrate-binding activity by pre-pro-conA in the endoplasmic reticulum and appears to be required for transport of pro-conA out of the endoplasmic reticulum (Faye and Chrispeels, 1987), and its removal is carried out by either an N-glycanase or endoglycosidase H in the protein body compartment of the seed (Sheldon and Bowles, 1992; Ramis et al., 2001). For brevity, we refer to deglycosylated pro-conA simply as pro-conA. In the protein body, an intervening 15-amino acid peptide (VIRNSTTIDFNAAYN) in the middle of the protein, where the glycan group was initially attached, is proteolytically excised by asparaginyl endopeptidase (AEP, sometimes referred to as vacuolar processing enzyme or legumain), creating new N-and C-termini. A new peptide bond is formed between the original N- and C-termini via a postulated cleavage-coupled transpeptidation event, wherein nine amino acids (EIPDIATVV) at the original C-terminus are removed (Bowles and Pappin, 1988) (**Figure 1B**). The excision of the 15-amino acid intervening peptide involves three cleavage reactions (N119, N130, N134); two of which (N119, N130) usually occurs before the transpeptidation event. The last cleavage reaction at N134 occurs much later after transpeptidation. This seemingly complex sequence of processing events resulting in the circular permutation of conA was at the time unprecedented (Carrington et al., 1985) and later was evocatively termed “protein carpentry” (Hendrix, 1991).

**Figure 1.**
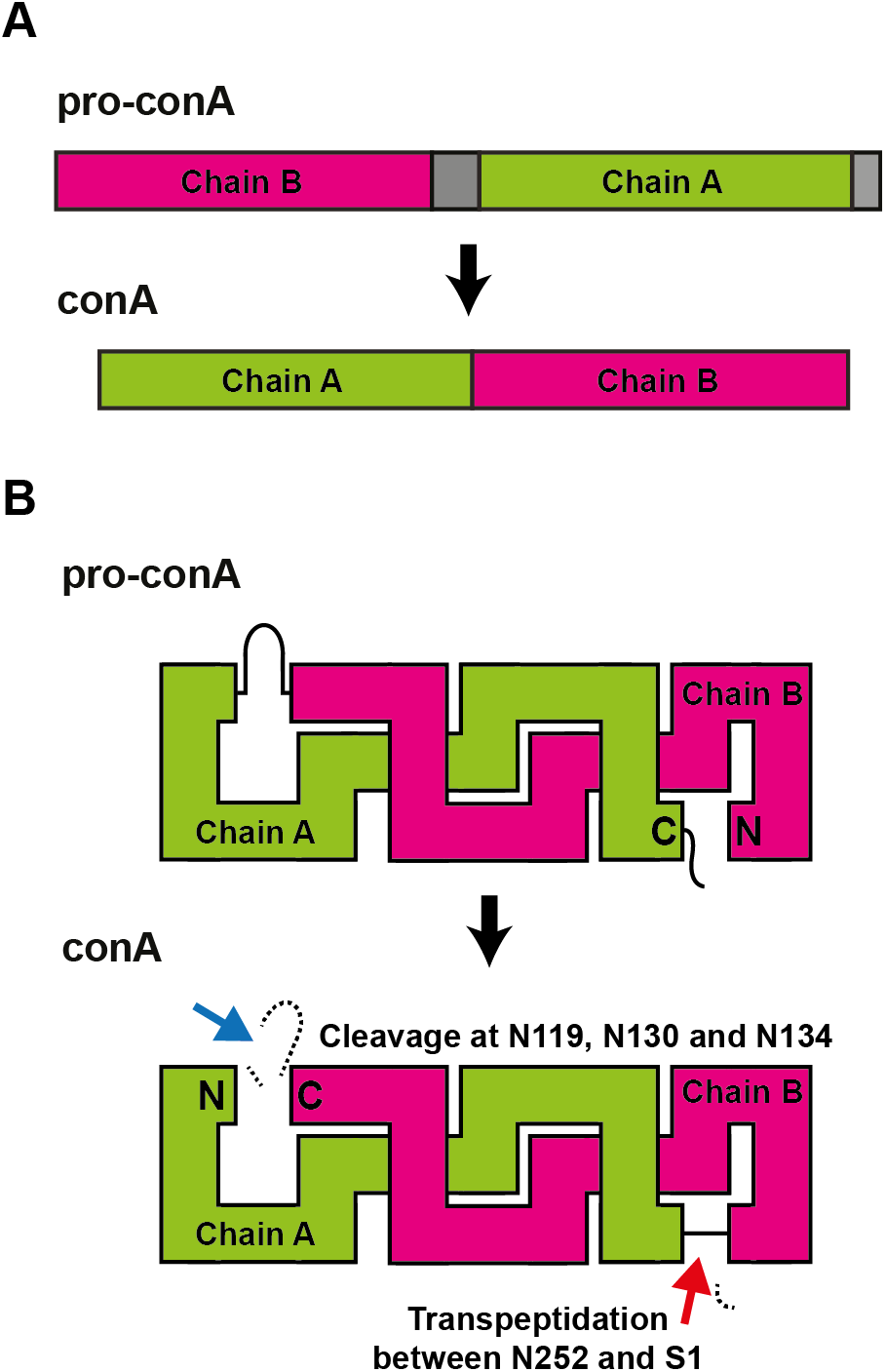
Concanavalin A maturation by circular permutation. (**A**) During maturation, the N-terminal half (Chain B) of pro-conA becomes the C-terminal half of conA and vice versa. Grey segments are cleaved off during maturation. (**B**) Peptide bond cleavages (blue arrow), and peptide bond formation i.e. transpeptidation (red arrow) within pro-conA are both required to make conA. The cleavage and transpeptidation events all occur on the carboxyl side of asparagine residues. The N-terminal ER signal of pre-pro-conA and the N-linked glycan, which are removed before circular permutation of pro-conA, are not shown.

Transpeptidation at the N- and C-termini appears to be an inefficient process as roughly half of conA in jack bean seeds exist in the two-chain form (Faye and Chrispeels, 1987), which is produced when the cleavage reaction is resolved by hydrolysis rather than aminolysis i.e. transpeptidation. The hydrolysed two-chain products are not degraded, which contrasts to what was observed in seeds of the common sunflower, where transpeptidation produce a macrocyclic peptide, and hydrolysis yields a small linear peptide that is degraded (Bernath-Levin et al., 2015). In solution, conA exists in a pH- and temperature-dependent, dimer-tetramer equilibrium (McKenzie et al., 1972; Huet and Claverie, 1978; Senear and Teller, 1981). ConA dimers consisting of a mixture of two-chain and full length protein are less competent at forming tetramers than purified conA consisting only of full-length protein (Senear and Teller, 1981). As conA in the dimeric and tetrameric forms appear to have different biological activities on animal cells *in vitro* (Gunther et al., 1973), it is possible that AEP-mediated processing of conA to produce both the two-chain and full length conA may be the result of as yet unknown selection pressures.

After the discovery that conA was circularly permuted, protein engineers began to modify proteins or enzymes in a similar way (Goldenberg and Creighton, 1983) to gain insights into protein folding (Gebhard et al., 2006) or to modify biophysical properties to overcome limitations (Meister et al., 2011; Yu and Lutz, 2011; Bliven and Prlić, 2012). Many more examples of circularly permuted proteins have since been discovered, but all of these natural examples, like the artificial circular permutations, are made at the genetic level. ConA and the closely-related conA-like lectins in the Diocleinae subtribe of plants remain the only proteins observed to undergo post-translational circular permutation (Cavada et al., 2018). The functional purpose for conA circular permutation remains elusive as few biochemical studies have been performed with purified pro-conA. Deglycosylation is the only modification necessary for pro-conA to acquire its carbohydrate-binding ability (Min et al., 1992; Sheldon and Bowles, 1992; Ramis et al., 2001). Furthermore, the cleaved, two-chain form of conA is capable of carbohydrate binding (Faye and Chrispeels, 1987), and homologs of pro-conA in other legumes do not undergo circular permutation, making it unclear why it occurs for conA (Cunningham et al., 1979; Carrington et al., 1985).

The maturation process of conA has been shown to involve a protease purified from jack bean seeds, which was characterised by Edman degradation of the first 25 residues (Abe et al., 1993; Min and Jones, 1994). This was the first enzyme discovered to be capable of forming peptide bonds post-translationally within the backbone of protein substrates (Min and Jones, 1994). Interest in exploiting this function for biotechnological applications, on top of efforts to identify similar enzymes capable of peptide backbone transpeptidation, has led to the discovery and engineering of AEPs with varying efficiencies in peptide backbone cleavage, transpeptidation and macrocyclisation, and AEPs with differing preferences for Asp and Asn residues (Nguyen et al., 2014; Yang et al., 2017; Haywood et al., 2018; Harris et al., 2019; James et al., 2019). However, our understanding of the AEP domains critical for determining hydrolase and transpeptidase efficiency is far from complete.

Although recombinant jack bean AEP (CeAEP1) has been shown to be capable of carrying out transpeptidation on non-native substrates *in vitro* (Bernath-Levin et al., 2015), it has not been shown beyond doubt to be capable of conA circular permutation. Here, we reconstitute the biosynthesis of conA using recombinant pro-conA and CeAEP1, with structural evidence supporting cleavage-mediated transpeptidation in conA. The vast majority of studies on conA have characterised its mature form. Here, we focus on why pro-conA undergoes circular permutation and the structural features of CeAEP1 that facilitate this reaction. We compare the carbohydrate binding ability and stability of pro-conA and conA by combining structural analyses with circular dichroism and isothermal titration calorimetry (ITC). Although circular permutation does not cause structural changes in the conA carbohydrate-binding domain, it results in increased thermal and chemical stability, which is consistent with fewer atomic interactions between subunits. The structure of the jack bean AEP is consistent with other related AEPs that carry out both cleavage and transpeptidation, explaining why only a single enzyme is required to circularly permutate pro-conA into conA.

## Materials and Methods

### Pro-conA expression and purification

A synthetic DNA sequence encoding residues 31-290 of *C. ensiformis* conA precursor (UniProt ID: P02866) was codon optimised for *E. coli* (GenScript) and included a Gly-Ser linker (encoded by an in-frame BamHI site) between a six-His tag and tobacco etch virus (TEV) protease recognition site (Glu-Asn-Leu-Tyr-Phe-Gln-Ser) at the N-terminus. Ser30 of UniProt ID: P02866 was not included because TEV cleavage will leave a serine residue at the N-terminus of pro-conA, hence reconstituting the native pro-conA protein sequence). Cultures were grown in lysogeny broth containing 100 μg/mL ampicillin and 35 μg/mL kanamycin at 30 °C and allowed to cool to 16 °C before inducing expression by adding 0.1 mM isopropyl β-D-1-thiogalactopyranoside at OD_600_ of 0.8-1.0. After 16 hours, cultures were harvested by centrifugation and lysed by ultrasonication in MOPS-salt buffer at pH 6.8 (50 mM MOPS (3-(N-morpholino) propanesulfonic acid), 12.5 mM sodium acetate, 1 M sodium chloride, pH 6.8). Cleavage with TEV protease was performed by first dialysing pro-conA into 50 mM sodium acetate, 500 mM sodium chloride, 1 mM dithiothreitol, pH 5.8 at 4 °C for 2 hours before centrifugation to remove precipitate that formed from the decrease in pH. His-tagged TEV protease was added (10% of pro-conA mass content) and the mixture incubated at 4 °C for a further 30 hours. To remove the TEV protease and the N-terminal tag cleaved from pro-conA, the mixture was dialysed overnight in MOPS-salt buffer then incubated (batch wise) with Ni-NTA resin overnight. Purified pro-conA was obtained in flow-through from reverse His-tag purification before a final concentration of 1 mM manganese chloride and 1 mM calcium chloride was added to purified pro-conA.

### Jack bean AEP expression and purification

A synthetic DNA sequence encoding residues 36-475 of *C. ensiformis* AEP (UniProt ID: P49046), including an N-terminal six-His tag and codon-optimised for *E. coli* (GeneArt), was sub-cloned into pQE30 (Qiagen) before being expressed in T7 Shuffle Express *E. coli* (New England Biolabs) containing the suppressor plasmid pREP4 (Qiagen). Expression was performed as above and cultures were harvested and lysed by ultrasonication in Tris-salt buffer (50 mM Tris pH 8.0, 100 mM sodium chloride) containing 0.1% Triton X-100. Lysed products were centrifuged and the supernatant incubated (batch wise) with Ni-NTA resin overnight at 4 °C. The resin was washed with 50 mL of Tris-salt buffer and 50 mL of Tris-salt buffer containing 20 mM imidazole before the recombinant protein was eluted with 20 mL of Tris-salt buffer containing 300 mM imidazole. Nickel-purified CeAEP1 was further purified by anion-exchange chromatography (HiTrap Q HP 5 mL) with gradient of 0 to 500 mM sodium chloride in 50 mM Tris, pH 8.0 over 90 min, followed by size exclusion chromatography (HiLoad 16/600 Superdex 200) in 50 mM Tris, 50 mM sodium chloride.

### Crystallisation and data collection

CeAEP1 was assessed for purity by SDS-PAGE. Low-resolution diffracting crystals were obtained initially using the sitting-drop vapour diffusion method with reservoir condition 100 mM sodium HEPES, 15% PEG 20,000 (w/v), pH 7 from the ProPlex crystallisation screen (Molecular Dimensions). An additive screen was then performed using the sitting-drop vapour diffusion method with 30 μL of reservoir solution (125 mM HEPES, 12.5% PEG 20,000 (w/v), pH 7.5) in a MiTeGen In Situ 1 crystallisation plate at 16 °C. Crystals were obtained from a condition with 0.5 μL of 16.7 mg/ml of protein, 0.4 μL reservoir solution, 0.1 μL additive (10 mM ethylenediaminetetraacetic acid disodium salt dihydrate) after 2 weeks. Single crystals were soaked in mother-liquor supplemented with 30% glycerol as a cryoprotectant prior to being flash frozen and stored in liquid nitrogen. Data collection was performed at 100 K on the Australian MX2 (micro-focus) beamline (McPhillips et al., 2002) using a wavelength of 0.9537 Å and diffraction data was collected to 2.7 Å resolution.

For pro-conA, a 10 kDa M.W.C.O. centrifugal filter (Amicon) was used to change buffer to 50 mM MOPS, 12.5 mM sodium acetate, 200 mM sodium chloride, 1 mM manganese chloride, 1 mM calcium chloride. Protein was assessed for purity by SDS-PAGE and concentrated to 28.7 mg/mL. Crystal screening was performed using the sitting-drop vapour diffusion method with 60 μL of reservoir solution in 96-well Intelli-Plates at 16 °C. Crystals were obtained from sitting drop containing 0.2 μL protein, 0.1 μL mother-liquor (10 mM zinc chloride, 100 mM HEPES, 20% PEG 6000 (w/v), pH 7.0) of the PACT Premier crystallisation screen (Molecular Dimensions). Single crystals were soaked in mother-liquor supplemented with 25% ethanediol as a cryoprotectant prior to being flash frozen and stored in liquid nitrogen. Data collection was performed similarly to CeAEP1 crystals and diffraction data was collected to 2.1 Å resolution.

### Structural determination, refinement and model building

For both pro-conA and CeAEP1, diffraction data were processed using XDS programme package (Kabsch, 2010) and scaled with AIMLESS from the CCP4 programme suite (Winn et al., 2011). Pro-conA crystallised with space group *I* 1 2 1 and unit cell dimensions a = 59.94 Å, b = 90.42 Å, c = 86.86 Å, β = 91.13°. The structure of pro-conA was solved by molecular replacement using MOLREP, with conA (PDB: 1JBC (Parkin et al., 1996)) as the search model. CeAEP1 crystallised with space group *I* 1 2 1 and unit cell dimensions a = 106.99 Å, b = 88.88 Å, c = 109.85 Å, β = 111.72°. The structure of CeAEP1 was solved by molecular replacement using MOLREP, with *Arabidopsis thaliana* legumain (PDB: 5NIJ (Zauner et al., 2018b)) as the search model. For both pro-conA and CeAEP1, manual building and refinement was performed in iterative cycles using COOT (Emsley et al., 2010) and REFMAC5 of the CCP4 programme suite. Coordinates and structure factors were deposited into the Protein Data Bank (PDB) under accession code 6XT6 (pro-conA) and 6XT5 (CeAEP1). Figures illustrating both structures were generated using PyMol. PyMol was used to calculate r. m. s. d. values. For comparison of pro-conA and conA functional sites, only the residues involved in the respective functions were aligned. CheckMyMetal (http://csgid.org/metal_sites) (Zheng et al., 2017) was used to evaluate the assignment of the metal binding sites in pro-conA.

### Confirming pro-conA maturation by CeAEP1 *in vitro*

Pro-conA maturation was performed by incubating 1 mg/mL of pro-conA with 0.1 mg/mL of CeAEP1 in MOPS storage buffer at pH 6.8 for 24 hours. Concentrations of pro-conA and conA were measured by absorbance at 280 nm with a NanoDrop, with protein extinction coefficient of 33920 M^−1^cm^−1^ and 32430 M^−1^cm^−1^, respectively, and molecular weight of 28.22 kDa and 25.57 kDa, respectively. Reaction was stopped by incubating in 5X sample loading buffer (20% (v/v) glycerol, 15% (w/v) sodium dodecyl sulfate, 312.5 mM Tris, 10 mM ethylenediaminetetraacetic acid disodium salt, 0.05% (v/v) β-mercaptoethanol, 0.05% (w/v) bromophenol blue, pH 6.9) at 37 °C for 15 minutes. Samples were run on SDS-PAGE containing 10 mM reduced glutathione in the running buffer. Samples were loaded to have roughly similar band intensity for the band of interest to enable better estimation of relative protein size (recombinant pro-conA: 0.75 μg, conA (Sigma): 1.5 μg, CeAEP1-processed conA: 6 μg). Samples were electrophoresed on a Bolt 4-12% Bis-Tris Plus gel, electroblotted onto Immobilon-PSQ PVDF membrane in Towbin buffer (25 mM Tris, 192 mM glycine, 10% (v/v) methanol) (Towbin et al., 1979) at 200 mA for 2 h in an ice water bath. Immobilon-PSQ PVDF membrane was then stained with 0.025% Coomassie Brilliant Blue R-250, 40% methanol. N-terminal Edman degradation of the first 8 amino acids was performed by Proteomics International (Perth, Australia).

### Circular dichroism

Pro-conA was concentrated to 10 mg/mL using 10 kDa M.W.C.O. centrifugal filter (Amicon) in MOPS-salt buffer. Concanavalin A purified from *C. ensiformis* (Type VI, lyophilised powder, Sigma-Aldrich, Cat No. L7647) was solubilised to 10 mg/mL in MOPS-salt buffer with 1 mM manganese chloride and 1 mM calcium chloride. Proteins were then diluted 100 times in water to 0.1 mg/mL, pH 6.5 immediately before CD melt curve analysis. Melt curves were measured at 218 nm with a temperature slope of 1 °C/min from 25-95 °C, 4-second response time, 5 nm bandwidth. A biphasic curve (GraphPad Prism, version 8.00) was fitted to the data. No-heat controls for pro-conA and conA were performed by measuring ellipticity at 25 °C for the same duration as the heat analysis. For pH stability analysis, pro-conA and conA were prepared in 20 mM Tris, pH5.8, 1 M sodium chloride, 1 mM calcium chloride, 1 mM manganese chloride before diluting 100 times for CD analysis. CD measurements were performed in triplicate using JASCO J-810 spectropolarimeter with quartz cuvette of 1 mm path length, 100 millidegree sensitivity, 1 nm data pitch, 100 nm/min scanning speed, 2 second response time, 4 nm bandwidth, 3 accumulations, between 190-260 nm at room temperature (24 °C).

### Isothermal titration calorimetry

Microcal iTC_200_ from GE healthcare was used to perform ITC analysis. Pro-concanavalin A (pro-conA) and Concanavalin A (conA), were dialysed in 50 mM MOPS, 12.5 mM sodium acetate, 1 M sodium chloride, 1 mM calcium chloride, 1 mM manganese chloride, pH 5.8, and concentrated down to 449.4 μM and 439.5 μM, respectively. The ligand, methyl-α-D-mannose, was dissolved to 8 mM with the same buffer used in protein sample dialysis. Protein samples were placed in the sample cell (cell volume = 200 μL) and titrated with methyl-α-D-mannose. Titrations were performed at 25 °C with a stirring speed of 750 rpm. Methyl-α-D-mannose was injected 76 times from a computer controlled syringe, at a volume of 0.5 μL over one second for each injection, with a spacing of 150 seconds between injections. Only 0.25 μL was injected for the first injection and ignored in analysis to minimise potential errors from preparation. Experimental data were fitted to a theoretical titration curve using the Origin® software (version 2002, OriginLab Corporation). A ‘one site model’ was used to generate the curve, with ∆H (enthalpy change), K_a_ (association constant) and the stoichiometry of the protein-ligand complex set as variable parameters.

## Results

### Pro-conA structure

To determine the structural differences between pro-conA and conA, we expressed recombinant pro-conA (residues 30-290; UniProt ID: P02866) in *E. coli* (**Supplemental Figure 2**). We obtained a protein crystal which diffracted to 2.1 Å by X-ray diffraction. The crystal structure was solved by molecular replacement using conA (PDB: 1JBC (Parkin et al., 1996)) as the search model, yielding a homodimer in the asymmetric unit. Superposition of the two monomers yields an r. m. s. d. of 0.4 Å over 225 Cα-atoms. Considerable chain mobility is inferred from poor electron density of the C terminus (residues 250-260) of both monomers. The C terminus is in close proximity to a solvent exposed loop (residues 65-70) which are involved in different crystal contacts in the two monomers. Residues at the functional sites discussed below are clearly visible in electron density (**Supplemental Figure 3**).

Like other lectins, pro-conA is made up of two large β-pleated sheets consisting of a flat six-stranded antiparallel β-sheet and a curved seven-stranded antiparallel β-sheet (**Figure 2A**). Pro-conA (chain A) and the conA monomer (PDB 1JBC) are very similar, with an r. m. s. d. of 0.5 Å over 221 Cα-atoms (**Supplemental Figure 4**). There is weak electron density for the 15-amino acid intervening peptide that gets cleaved off during conA maturation. Crystals of pro-conA were therefore dissolved and run on an SDS-PAGE gel to confirm presence of full-length protein, demonstrating that weak electron density for the 15-amino acid intervening peptide is due to disorder rather than spurious cleavage by other enzymes during expression or purification (**Supplemental Figure 2**).

**Figure 2.**
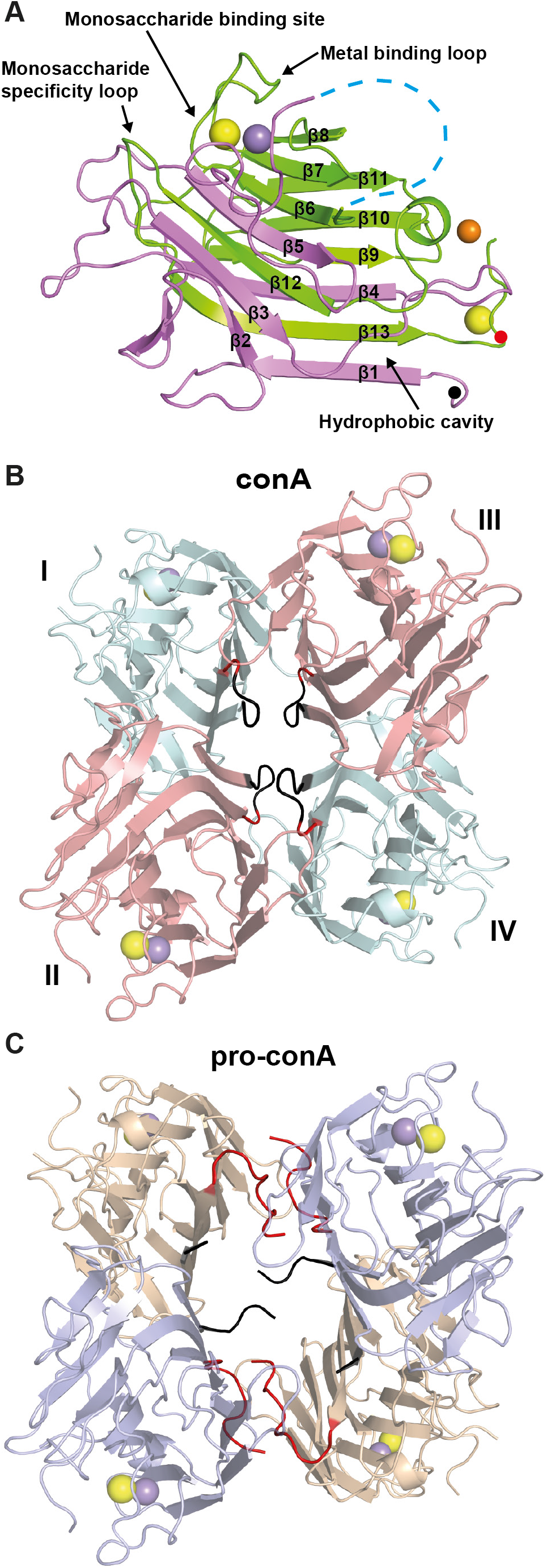
Pro-conA and conA have similar monomeric structures but different tetrameric structures in protein crystals. (**A**) Pro-conA consists of interleaved β-sheets (β1-β13) from chain A (pink) and chain B (green). Transpeptidation occurs between the N-terminus (black dot) and a residue near the C-terminus (red dot). ‘Metal binding loop’ and ‘monosaccharide specificity loop’ are a single loop each whereas the ‘monosaccharide binding site’ consists of residues from three loops. The ‘hydrophobic cavity’ consists of residues from β1, β13 and a nearby loop. Fifteen flexible residues linking chains A and B (blue dashed line); ions of calcium (yellow sphere); manganese (purple sphere); zinc (orange sphere). This tertiary structure is similar to conA (**Supplemental Figure 3**). (**B**) In conA, monomers I and II interact so that the flat six-stranded β-sheet from each monomer aligns to form a contiguous 12-stranded β-sheet. Monomers III and IV dimerises in a similar manner. The two dimers then interact via the 12-stranded β-sheets to form a dimer-of-dimers. (**C**) Pro-conA adopts a dimer-of-dimers conformation with a larger cavity. The pro-conA termini interfere with the formation of the typical dimer-of-dimers. N- and C-termini of pro-conA monomers are shown in black and red, respectively. The equivalent residues in conA are coloured similarly.

### Structural features involved in conA activity

It is not known if conA circular permutation changes the structural domains involved in carbohydrate binding. Here we see that the monosaccharide binding site of pro-conA and conA are very similar, with an r.m.s.d. of 0.2 Å for the three residues involved (**Supplemental Figure 3A**). The monosaccharide specificity loop of pro-conA and conA have an r.m.s.d. of 1.1 Å for the six residues involved (**Supplemental Figure 3B**). Differences in the sidechain positions of Leu233 and Tyr234 of the monosaccharide specificity loop, which has a nearby ethanediol molecule in the pro-conA structure, are unlikely to affect carbohydrate binding as only their main chain atoms interact with carbohydrates (Kanellopoulos et al., 1996b; Hamodrakas et al., 1997). Other residues involved in carbohydrate recognition, including Tyr146-Ile151 (Tyr12-Ile17 in conA) and Thr108-Leu111 (Thr226-Leu229 in conA) (Loris et al., 1998; Cavada et al., 2018), are also structurally unchanged after circular permutation.

ConA carbohydrate-binding activity is highly dependent on the binding of a transition metal, typically manganese, and a calcium ion (Sumner and Howell, 1936; Kalb and Levitzki, 1968; Shoham et al., 1973). The presence of transition metal ions and calcium has also been shown to improve conA structural stability (Blumberg and Tal, 1976; Doyle et al., 1976). In this structure, a manganese ion is coordinated by Glu142, Asp144, Asp153, His158 (Glu8, Asp10, Asp19, His24 in conA) and two water molecules, and a calcium ion is coordinated by Asp144, backbone of Tyr146, Asn148, Asp153 (Asp10, Tyr12, Asn14, Asp19 in conA) and two water molecules. The binding of these two metals has been shown to stabilise the active ‘locked’ conformation in conA (Brown et al., 1977; Brewer et al., 1983; Bouckaert et al., 2000), which is observed here in metallated pro-conA as well (**Supplemental Figure 3C)**. As proposed in the conA structure by Bouckaert and colleagues, in pro-conA the coordination of Asp144 and Tyr146 (Asp10 and Tyr12 in conA) to the calcium ion induces a bend in the β-sheet holding Thr145 (Thr11 in conA), causing a steric clash between Thr145 and Asp90 (Asp208 on conA) and inducing a trans-to-cis isomerization between Ala89 and Asp90 (Ala207 and Asp208 in conA). The key residues indicating a locked conformation in pro-conA is similar to those of conA, with an r.m.s.d. of 0.8 Å for the five key residues highlighted in **Supplemental Figure 3c**.

There is a small possibility that the transition-metal-binding site may be occupied by a nickel ion rather than a manganese ion. Although we cannot rule out the presence of nickel in the model, as nickel was used in pro-conA purification, the overall geometry of the metal binding sites have been shown to be essentially independent of the nature of the transition metal (Emmerich et al., 1994). Zinc and calcium ions were modelled at four other locations in the homodimer and are not part of the metal binding loop. There is potentially a mixture of metals in these locations, but these metals are unlikely to have a biological role and are likely present due to crystallisation conditions.

A hydrophobic cavity conserved in conA-like lectins has been hypothesised by some to bind to secondary metabolites (Delatorre et al., 2007; Bezerra et al., 2011). An ethanediol (cryoprotectant) is observed in this cavity in the pro-conA structure, and the local structure appears unchanged when compared to a conA structure containing ethanediol in this position (PDB:4PF5 (François-Heude et al., 2015)), with an r.m.s.d. of 0.2 Å over three residues **Supplemental Figure 3D**).

### Pro-conA and conA interact differently in protein crystals

In contrast to the well-established dimer-of-dimers complex (i.e. tetramer) observed in the conA crystal structure (**Figure 2B**), pro-conA monomers in the crystal structure assemble in an atypical dimer-of-dimers complex (**Figure 2C**). The asymmetric unit contains a dimer, composed of very similar interactions as observed in the principal dimer of conA (involving residues 3-21, 57-65, 99, 222-224 and 248-252 in pro-conA). Application of crystal symmetry to the coordinates yields a possible tetrameric structure. PDBePISA (Krissinel and Henrick, 2007) was used to predict the free energy of assembly-dissociation (ΔG^diss^) for pro-conA, indicating that pro-conA is likely capable of forming a stable dimer (ΔG^diss^ = 5.0 kcal/mol) and tetramer (ΔG^diss^ = 17.2 kcal/mol) in solution. Positive values of ΔG^diss^ indicate that an external driving force should be applied to dissociate the assembly, therefore assemblies with ΔG^diss^>0 are thermodynamically stable. For comparison, high resolution structures of conA (e.g. PDB: 1JBC) also predict a stable dimer (ΔG^diss^ = 8.6 kcal/mol) and tetramer (ΔG^diss^ = 9.6 kcal/mol) in solution. Compared to conA, however, pro-conA has a smaller buried surface area and fewer interface residues, especially between the dimers (**Supplemental Table 1**). PDBsum (Laskowski et al., 2018) was used to analyse the number of non-covalent contacts with a distance cut-off of 4 Å, and this showed that pro-conA has fewer inter-dimer contacts than conA in the crystal structure (**Figure 3**). As shown in **Figure 2C**, **3A** and **3C**, the C-termini of the pro-conA subunits are wedged between the dimers and are therefore heavily involved in the inter-dimer interactions. These termini are cleaved off and shortened into a loop during circular permutation to form conA. Although the region spanning four β-sheets (residues 183-210) is involved in inter-dimer interactions in both pro-conA and conA, they have different interacting partners. The residues in this β-sheet region in pro-conA interact with the wedged C-termini, while in conA they interact with the β-sheet region of the opposing subunit (**Figure 3**). Pro-conA also has substantially fewer interactions spanning residues 69-78.

**Figure 3.**
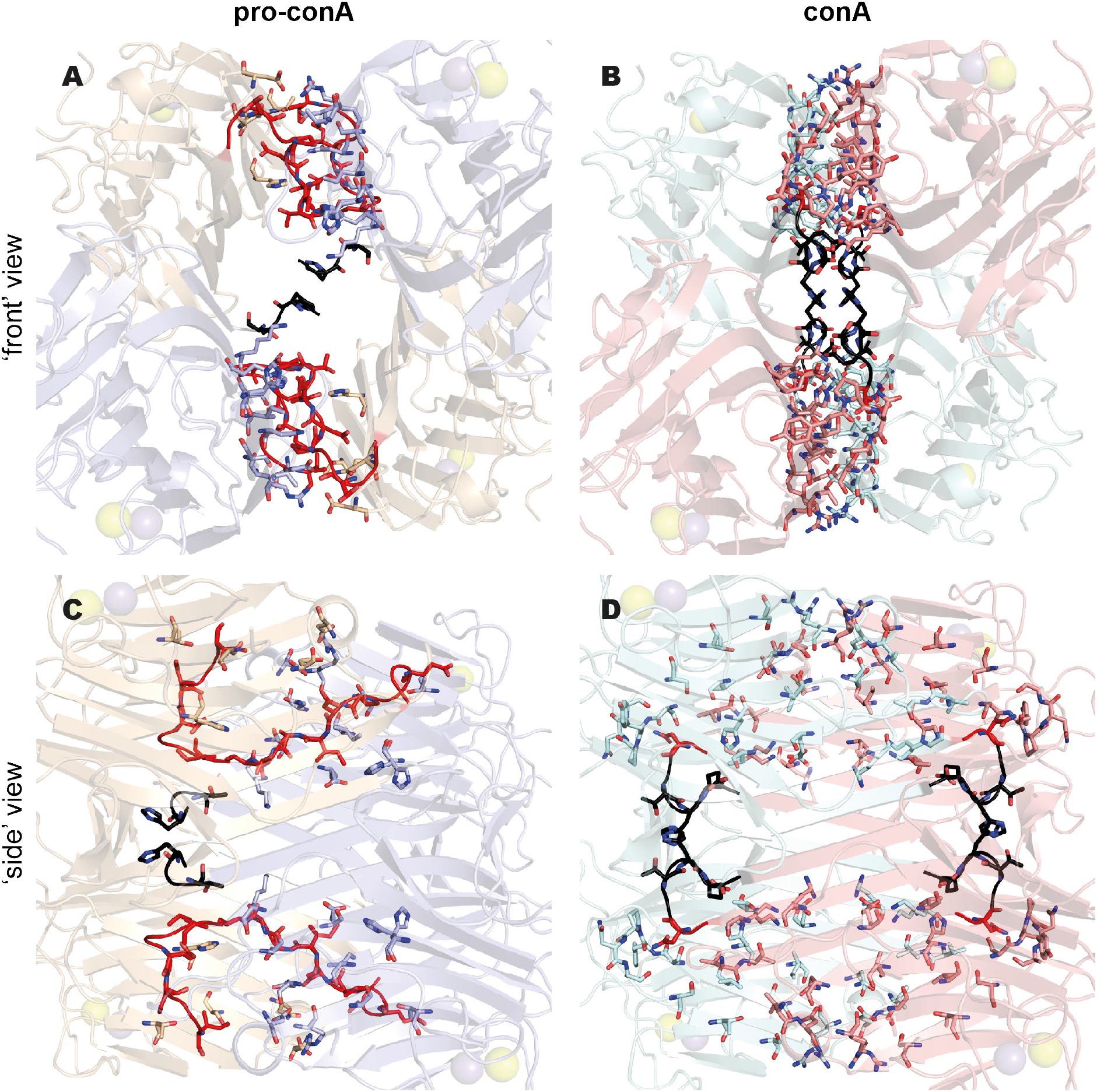
The pro-conA tetramer has fewer inter-dimer contacts than the conA tetramer in crystal structures. Pro-conA (**A** and **C**) has 94 non-bonded contacts between dimers in a dimer-of-dimers complex. ConA (**B** and **D**) has 194 non-bonded contacts between dimers in a dimer-of-dimers complex. In pro-conA, the C-termini interfere with the inter-dimer interactions that would exist between the opposing β-sheets of conA. Residues involved in non-bonded contacts are shown as sticks. The ‘front’ view shows all four subunits of the dimer-of-dimers. ‘Side’ view is obtained with a 90° anti-clockwise turn when viewed from the top. The N- and C-termini are highlighted in black and red respectively.

### Reconstituting conA biosynthesis *in vitro*

To determine whether CeAEP1 alone can perform separate cleavage and cleavage-coupled transpeptidation reactions to make conA, we sought to reconstitute conA biosynthesis *in vitro* with recombinant pro-conA. Recombinant pro-conA (residue 30-290 of UniProt ID: P02866) was incubated with CeAEP1 at a molar ratio of 10:1 pro-conA:CeAEP1. The pro-conA band was converted to several lower MW proteins after digestion (**Figure 4A**). The product of interest in the lane of the pro-conA digest was the top-most band, which has a slightly smaller MW than pro-conA, consistent with a circularly permuted conA. This protein band was sequenced by N-terminal Edman degradation and this yielded two N-terminal sequences; AAYNADTI and ADTIVAVE. These overlapping sequences obtained by Edman degradation both prove that the C-terminal half of conA has been circularly permuted to become the N-terminal half, and concurs with the slightly larger MW observed for band I (**Figure 4**; **Supplemental Figure 5**). In jack bean seeds, the removal of the AAYN tetrapeptide preceding the N-terminus of conA occurs very slowly (Bowles et al., 1986), explaining why in this *in vitro* experiment we observed both species. This AAYN tetrapeptide was also observed in conA extracted from immature jack bean seeds (Wang et al., 1971; Carrington et al., 1985). More than half of pro-conA appears to undergo cleavage rather than transpeptidation at the termini (**Figure 4A**). This inefficient transpeptidation reaction was also observed in conA extracts from jack bean seeds (Carrington et al., 1985). The bottom three MW bands represent the cleaved products (Sheldon et al., 1996) (**Supplemental Figure 1**).

**Figure 4.**
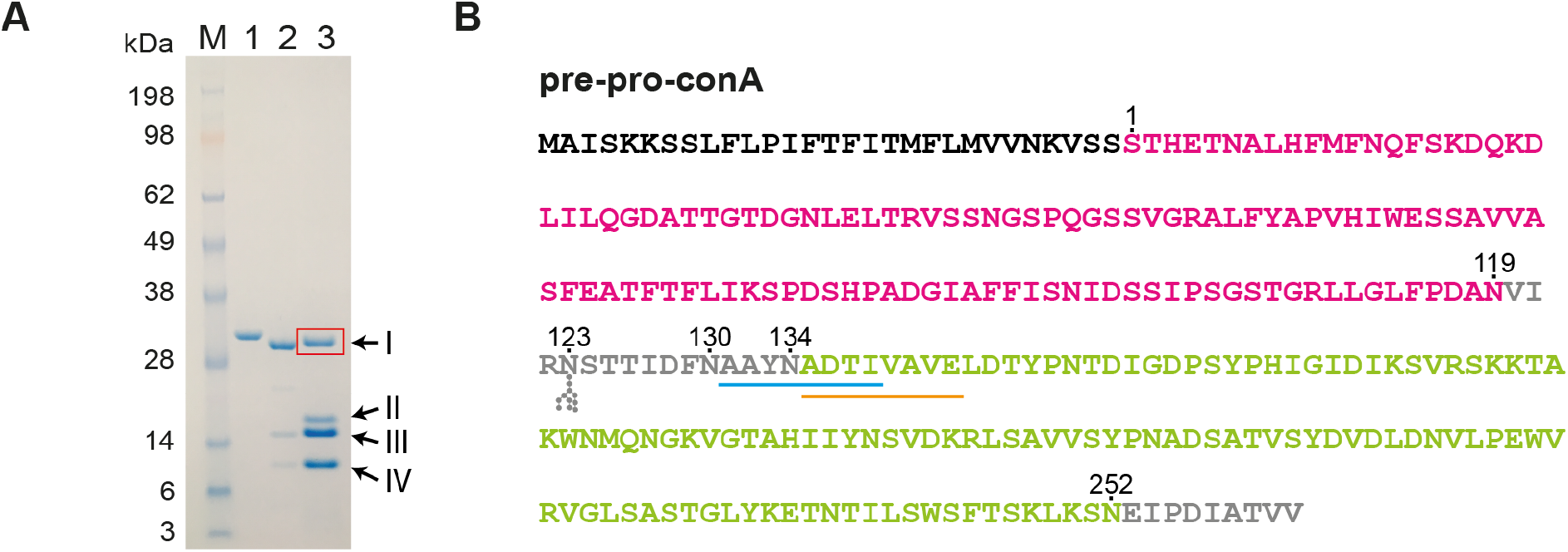
*In vitro* reconstitution of conA maturation using recombinant CeAEP1 and pro-conA. (**A**) SDS-PAGE of recombinant pro-conA (Lane 1), conA (Sigma-Aldrich, Cat No. L7647; Lane 2), and pro-conA processed by CeAEP1 (Lane 3). N-terminal sequence of Band I (red box) was determined by Edman degradation. The slightly larger size than conA is due to the presence of the AAYN tetrapeptide, which is cleaved off at a slow rate, on the N-terminus of circularly permuted conA (**Supplemental Figure 1**). Band III and Band IV are present in jack bean extract as well (Lane 2) and represent fragments that have undergone cleavage instead of transpeptidation (Sheldon et al., 1996). Band II likely represents cleaved fragment containing N-terminal AAYN tetrapeptide and/or the C-terminal EIPDIATVV peptide. (**B**) Pre-pro-conA sequence with ER signal (black), chain B (pink), chain A (green), and residues removed during circular permutation (grey). To facilitate comparison with pro-conA structure and previous studies, residue numbering starts after N-terminal ER-signal, at S1. Pro-conA is synthesised with a glycosylated N123 residue. Deglycosylation is the first post-translational modification. Cleavage by CeAEP1 occurs at N119, N130 and N134. N252 is either cleaved or transpeptidated to S1. The two N-terminal sequences from Edman degradation are underlined in blue and orange.

### Thermal and pH stability of pro-conA and conA

Circular dichroism analysis showed that conA is more stable than pro-conA under heat stress at pH 6.5 and at various other pH conditions without heat stress, revealing a functional consequence of conA circular permutation. During heat stress at pH 6.5, where temperature was increased at a rate of 1 °C /min, there was an increase in the magnitude of ellipticity at 218 nm for pro-conA at the 60-70 °C range before decreasing from 70 °C onwards (**Figure 5A**). In contrast, only a decrease in magnitude of ellipticity was observed for conA. This suggests that, under heat stress, pro-onA undergoes an increase in the amount of β-structure relative to conA. Controls showed no change in ellipticity for pro-conA and conA when no heat is applied **Supplemental Figure 6A**). SDS-PAGE analysis was carried out to illustrate the precipitation observed for both pro-conA and conA during the heat analysis (**Supplemental Figure 6B**). A previous study combining CD analysis with scattered light intensity analysis showed that conA begins to aggregate before any conformational change occurs during heat stress (Maeda et al., 1989), which may explain why we do not see an increase in magnitude of ellipticity for conA in our CD analysis. Even though pro-conA and conA appear to react differently to heat stress, it is clear that conA has better heat tolerance as pro-conA begins to undergo changes in conformation and/or begins precipitating at a lower temperature than conA at pH 6.5, and also appears to approach complete precipitation at a lower temperature than conA (**Figure 5A**). The melt curve for conA agrees with a previous study that observed conA aggregation from 60 °C onwards (Doyle et al., 1976).

**Figure 5.**
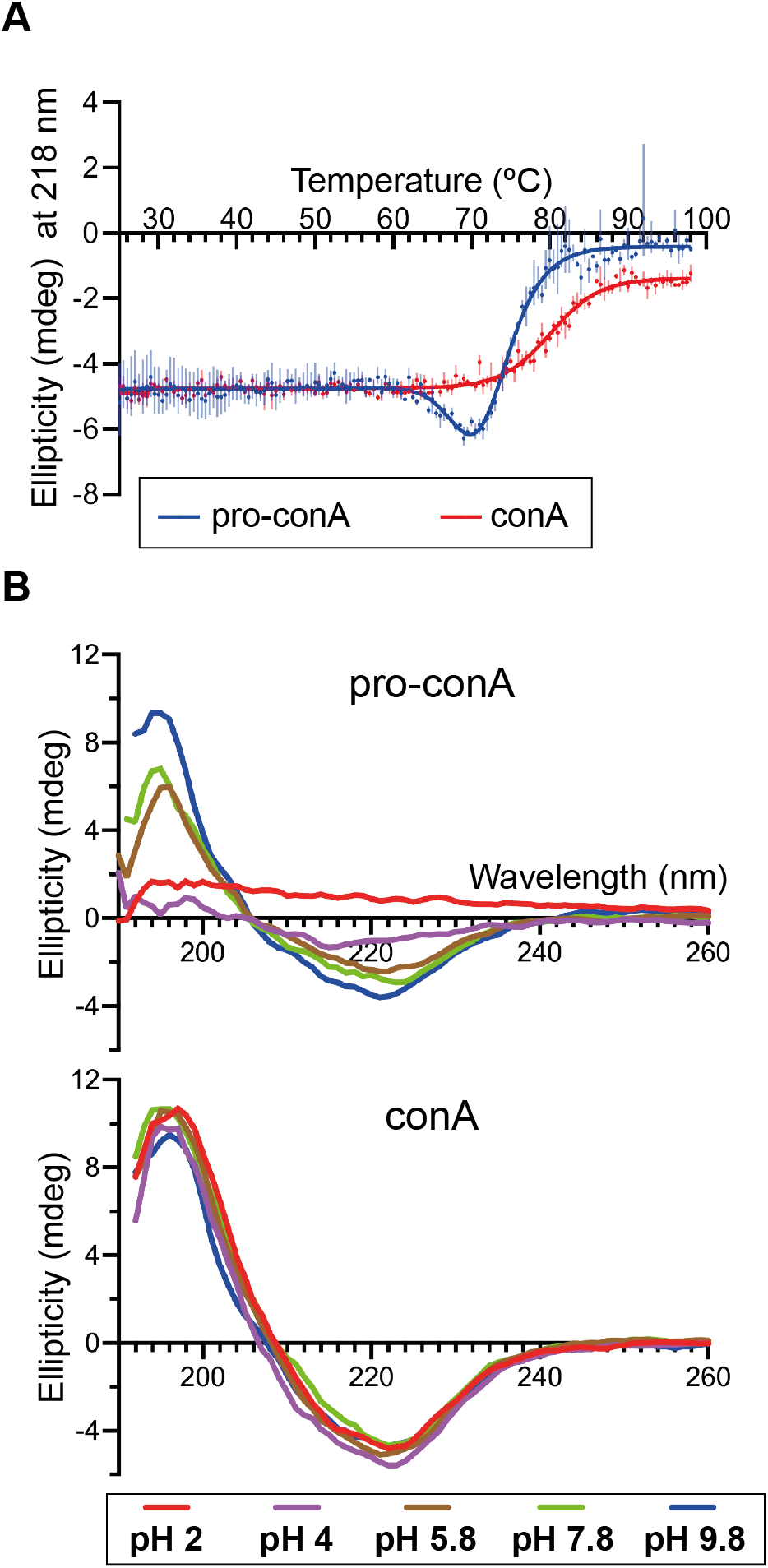
ConA is more stable than pro-conA. (**A**) Melt curve generated from circular dichroism analysis of pro-conA and conA in water with a temperature slope of 1 °C/min. Pro-conA begins changing conformation and precipitates from solution (**Supplemental Figure 8B**) at a lower temperature than conA. The decrease in ellipticity for pro-conA but not conA indicates that these two proteins have different pathways of unfolding and/or aggregation. (**B**) Circular dichroism spectra of pro-conA and conA after 2-hour incubation in water at pH 2, 4, 5.7, 7.8, 9.8. ConA remains stable from pH 2 to pH 9.8 whereas pro-conA is less stable at neutral and low pH.

To examine pH stability, pro-conA and conA were incubated at various pH conditions (pH 2 to 9.8) for two hours before CD analysis at the respective pH. CD spectra analysis showed that conA maintains its general structure throughout the pH range tested for the two-hour period of the analysis, whereas pro-conA is more prone to precipitation at lower pH conditions (**Figure 5B**). Precipitation occurred similarly to what was observed during heat stress (data not shown). Beyond the two-hour time frame, conA undergoes pH-dependent conformational changes as well (McCubbin et al., 1971; Zand et al., 1971).

The high similarity of the recombinant pro-conA structure that we obtained to that of conA indicates that recombinant pro-conA folds into the correct conformation in our *E*. *coli* expression system. The different sources for pro-conA and conA, *E. coli* and jack bean, respectively, are therefore an unlikely contributing factor to the differences observed in the biophysical assays. It should be noted that conA (Sigma) contains a small, unquantified amount of the two-chain form (**Figure 4A**), representing the products that have undergone cleavage rather than transpeptidation.

### Circular permutation has no effect on conA binding to methyl-α-D-mannose

To determine if conA circular permutation affects carbohydrate binding, we investigated the ability for pro-conA and conA to bind to mannose, the preferred ligand *in vitro*, using ITC. The results were similar for pro-conA and conA, with association constants (K_A_) of 8270±74.0 M^−1^ and 8240±74.0 M^−1^, respectively, for methyl-α-D-mannose (**Supplemental Figure 7**). The K_A_ we obtained for conA was in close agreement with that obtained by Loka *et al.* (2015) and Chervenak *et al.* (1995), with both studies obtaining a K_A_ of 7.6 × 10^3^ M^−1^ for methyl-α-D-mannose.

### Jack bean AEP Structure

To understand the enzymatic basis for conA maturation, we expressed CeAEP1 (residues 36-473; UniProt ID: P49046) in *E. coli* (**Supplemental Figure 8**). We obtained a protein crystal which diffracts to 2.7 Å by X-ray diffraction. The crystal structure was solved by molecular replacement using *Arabidopsis thaliana* legumain γ (AtLegγ) as the search model, yielding a CeAEP1 homodimer in the asymmetric unit. The superposition of the two monomers yields an r. m. s. d. of 0.4 Å over 424 Cα-atoms. Residues at the catalytic site and the substrate binding pockets are clearly visible in electron density (**Supplementary Figure 9A** **and** **9B**). Differences between monomers are only at solvent-exposed residues in or adjacent to loop regions. Electron density shows clear differences at residues 283-289 and 347-360, which can be attributed to different crystal contacts. Electron density shows static disorder at residues 318-330, which contains the linker region that is cleaved off upon CeAEP1 autoactivation. Missing electron density at residues 323-329 when contoured at 1σ point to the highly flexible nature of this linker.

CeAEP1 is structurally similar to previously solved plant AEP structures: *Helianthus annuus* AEP1 or HaAEP1 6AZT (Haywood et al., 2018), AtLegγ 5NIJ (Zauner et al., 2018b), *Oldenlandia affinis* AEP1 or OaAEP1 5H0I (Yang et al., 2017), *Viola yedoensis* peptide asparaginyl ligase 2 or VyPAL2 6IDV (Hemu et al., 2019), and butelase1 6DHI (James et al., 2019) with an r. m. s. d. of 1.0-1.1 Å over 400-430 Cα-atoms between CeAEP1 and the other published structures (**Supplemental Table 2**). The zymogen is made up of a ‘core’ domain (Glu38-Asp313) linked to a C-terminal ‘cap’ domain (Arg336-Ala474) via the flexible linker, which has weak electron density (**Figure 6A**). The core domain consists of a six-stranded β-sheet surrounded by five major α-helices, and is highly structurally conserved with an r. m. s. d. of 0.5-0.7 Å over 270-280 Cα-atoms between CeAEP1 and the other published structures (**Supplemental Table 3**). The cap domain, consisting of five α-helices, improves core domain stability, acts as the dimer interface and modulates enzymatic activity by occluding the active site in the zymogenic form (Zauner et al., 2018b).

**Figure 6.**
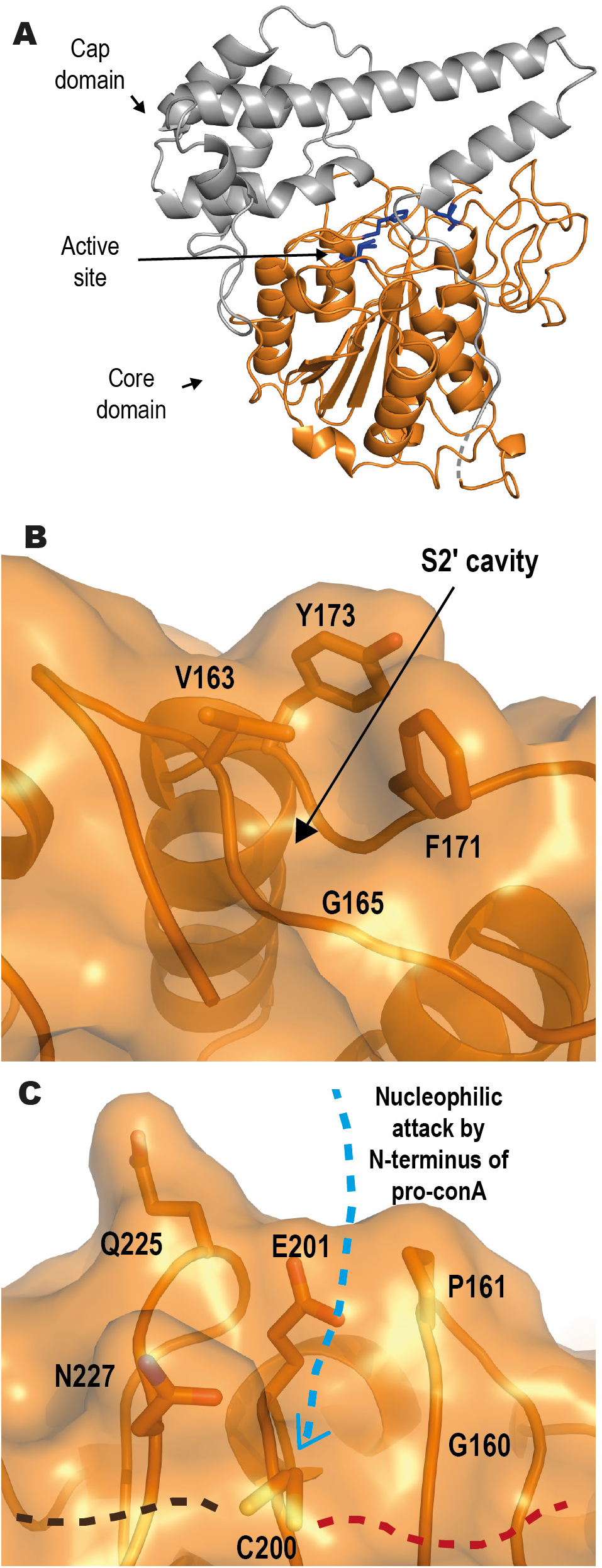
Jack bean asparaginyl endopeptidase 1 (CeAEP1) structure. (**A**) Inactive CeAEP1. Core domain (orange); cap domain; Active site (blue residues); Low electron density in linker region (dashed grey). (**B**) Gly165 in S2′ pocket forms a deep cavity, possibly facilitating the binding of peptide substrate and preventing replacement with water molecules. (**C**) Residues G160, P161, E201, Q225 and N227 likely interfere with incoming N-terminus of peptide substrate during nucleophilic attack, resulting in poor transpeptidase efficiency. C200 is the catalytic residue. Incoming N-terminus of pro-conA with nucleophilic serine (blue dashed arrow) attacking the acyl-enzyme intermediate at the catalytic cysteine; Residues N-terminal to cleavage site on pro-conA (brown dashed line); Residues C-terminal to cleavage site of pro-conA (red dashed line).

### CeAEP1 active site residues are strictly conserved

The catalytic residues His158 and Cys200 form the catalytic dyad, with Asn53 sometimes included to make up a catalytic triad (**Supplemental Figure 9A**). These three residues are conserved across all AEPs (**Supplemental Figure 10**). The His158 imidazole deprotonates Cys200, which becomes more reactive and catalyses the enzymatic activity. Electron density at Cys200 indicates a dual conformation, which is also observed in the near-atomic resolution structure of a HaAEP1, likely representing conformational flexibility that can occur during catalysis (Haywood et al., 2018). Mutagenesis study on HaAEP1 shows that the asparagine residue of the catalytic triad (Asn53 in CeAEP1; Asn73 in HaAEP1) influences the ratio of cyclic to acyclic product, but is not a strict requirement for cleavage or transpeptidation (Haywood et al., 2018). We modelled a succinimide intermediate (SNN) at Asp157 to obtain a better fit with the electron density at this position (**Supplemental Figure 9A**). The presence of SNN is consistent with all other plant structures except OaAEP1 (Yang et al., 2017; Haywood et al., 2018; Zauner et al., 2018b; Hemu et al., 2019; James et al., 2019), although OaAEP1 does have electron density suggesting that succinimide may in fact be present (James et al., 2019). Gln335, which is part of the cap domain, occupies the S1 pocket of the active site in the zymogen (according to nomenclature by Schechter and Berger where P1, P2 and so on refer to residues N-terminal to the cleavage site, whereas P1′, P2′ and so on are C-terminal to it, and where the corresponding binding sites on the protease are termed S2, S1, S1′, S2′, etc.) (Schechter and Berger, 1967) (**Supplemental Figure 9A**). It was hypothesised that cleavage does not occur at Gln335 even though it is similar to asparagine because the glutamine backbone is kept further away from the catalytic cysteine by virtue of the longer side chain (4.9 Å in CeAEP1) (Zauner et al., 2018a).

### CeAEP1 substrate binding pockets

With such high structural similarities amongst plant AEPs, CeAEP1 catalytic preference and efficiency can be explained only by subtle variations in the substrate-binding pockets. The S1 pocket (Arg55, His56, Glu198, Ser228, and Asp250), as designated by a previous study on AtLegγ (Zauner et al., 2018a), is conserved (**Supplemental Figure 9C**) and accommodates the P1 asparagine residue at all pro-conA cleavage and transpeptidation sites. His158, Gly159, Gly160, Cys200 and Glu201 make up the S1′ pocket (Zauner et al., 2018a; Hemu et al., 2019) and is highly conserved with notable variability in side chain positioning of Glu201 (**Supplemental Figure 9D**). Val163, Gly165, Phe171 and Tyr173 make up the S2′ pocket, with only Gly165 strictly conserved (**Supplemental Figure 9E**). The Phe171 equivalent in HaAEP1 (His191) adopts a different conformation from the corresponding residues in the other five AEP structures. This may be due to the interaction with a nearby glycerol molecule, or an effect of the charged histidine side chain as opposed to the usual hydrophobic side chain in this location in the other AEPs. The vast majority of the residues in the S2′ pocket in CeAEP1 and other AEPs are otherwise hydrophobic. The Arg278-Met294 region, which was designated the Marker of Ligase Activity by Jackson et al. (2018), is generally hydrophilic in CeAEP1, similar to that of HaAEP1 (**Supplemental Figure 10**).

## Discussion

It is just over 100 years since the discovery of conA. Despite conA being the subject of many studies and seeing daily use in laboratories worldwide, only now do we have a structure for pro-conA; this allows us for the first time to understand how circular permutation of pro-conA affects this long-studied protein.

### Circular permutation affects conA crystal packing

The structure of pro-conA confirms the long-standing suspicion that circular permutation of conA occurs without any large changes in the three-dimensional structure (Carrington et al., 1985) (**Supplemental Figure 4**). We observed a difference, however, in the number of inter-dimer interactions between the pro-conA and conA crystal structures (**Figure 3**; **Supplemental Table 2**). Even though conA in the de-metallated and the saccharide-bound form have different crystal packings, both forms adopt the typical dimer-of-dimers assembly (Kanellopoulos et al., 1996a). In pro-conA, the C-terminus is heavily involved in inter-dimer interactions (**Figure 2C**; **Figure 3A** **and** **C**). During circular permutation, the N- and C-termini undergo cleavage-coupled transpeptidation and are therefore present as a shortened loop that does not interfere with the formation of the typical dimer-of-dimers assembly in the conA crystal structure (**Figure 2B**; **Figure 3B** **and** **D**). It is likely that the inability for pro-conA to form the typical dimer-of-dimers assembly in the crystal structure is attributable solely to the presence of the termini. Glycinin, a seed storage protein in soybean, exists as a trimer before cleavage at an asparagine residue by an AEP triggers hexamer formation (Jung et al., 1998; Adachi et al., 2001; Adachi et al., 2003). Even though glycinin and conA are structurally different, in both cases AEP processing results in the dislocation of a peptide chain that appears to interfere with the formation of a higher oligomeric state. It would be interesting to see if the typical dimer-of-dimers conformation forms if pro-conA termini underwent AEP-mediated cleavage rather than transpeptidation.

### Binding domains in pro-conA remain unchanged after circular permutation

The requirement for conA to be in the ‘locked’ conformation to effectively bind carbohydrates has been explained in detail by in-depth structural analyses by Bouckaert et al. (2000). Here we obtain a structure of pro-conA which is similar to the ‘locked’ conformation (**Supplemental Figure 3C**), which explains why circular permutation is not required for carbohydrate binding and that the binding of Mn^2+^ and Ca^2+^, even in pro-conA, is sufficient for carbohydrate-binding activity (Ramis et al., 2001). This was corroborated with ITC analysis showing no difference in association constant between pro-conA and conA when assayed with methyl-α-D-mannose. The similar conformation of both the monosaccharide specificity loop and the monosaccharide binding site between pro-conA and conA indicates that circular permutation does not affect carbohydrate specificity either. A conserved hydrophobic cavity, hypothesised by Bezerra et al (2011) and Delatorre et al. (2007) to play a role in plant defence in their study of conA-like lectins, also remains unchanged by circular permutation.

### Circular permutation improves conA stability

Artificial circular permutation has on only one occasion led to a slightly more stable version of protein (Topell et al., 1999); this explorative modification otherwise tends to result in proteins that are either less or equally stable as the parent protein (Heinemann and Hahn, 1995). Here, we discover a clear improvement in stability resulting from protein circular permutation. We demonstrate that conA circular permutation increases conA thermal and pH stability (**Figure 5**) without affecting *in vitro* binding to the carbohydrate methyl-α-D-mannose (**Supplemental Figure 7**). The exceptional pH stability of conA was indeed a useful property for its initial discovery, as conA was readily separated from other seed proteins using a protocol involving a range of pH (Sumner, 1919). A more comprehensive binding analysis with various carbohydrates of different oligomeric forms may be useful, but the carbohydrate-binding function of conA *in vivo* has so far only been supported by *in vitro* characterisation rather than direct functional evidence in jack bean plants.

### CeAEP1 prefers serine over alanine as nucleophile in conA circular permutation

AEP cleavage at the pro-conA intervening peptide at Asn130 and Asn134 generates new alanine N-termini (Ala131 and Ala135) which do not undergo transpeptidation with Asn119 despite their close proximity. This selectivity was also observed in Sunflower Trypsin Inhibitor-1 (SFTI-1), where transpeptidation occurs when the incoming N-terminal nucleophile in native SFTI-1 is glycine, but a Gly1 to Ala1 mutation inhibits transpeptidation (Bernath-Levin et al., 2015). *In silico* docking simulations performed by Zauner et al. (2018a) indicate that an ionic interaction may facilitate transpeptidation between the N-terminal nucleophile and Glu220 of AtLegγ (Glu201 in CeAEP1) in the S1′ pocket (**Supplemental Figure 9D**). Transpeptidation in pro-conA occurs with an incoming N-terminal serine, which is similar in size to alanine. It is therefore possible that the polar sidechain of serine facilitates transpeptidation by CeAEP1 via ionic interactions, whereas the hydrophobic, non-polar property of Ala131 and Ala135 in pro-conA hinders the formation of this ionic interaction, thereby preventing transpeptidation by CeAEP1.

### CeAEP1 is predominantly a protease

The jack bean AEP structure not only provides a structural image of the enzyme responsible for conA circular permutation, but also improves our understanding on AEPs, a family of enzymes that are becoming recognised as a tool for protein engineering (Hemu et al., 2016; Yang et al., 2017; Tang et al., 2020). The structural features that make AEP favour hydrolysis or transpeptidation remains a matter of debate. Recent work by Du *et al.* (2020) uncovered a plant AEP with one of the highest transpeptidase activities despite predictions, based on previously published AEPs, that it would be a hydrolase. Here we compared CeAEP1 structure to butelase1, an extremely efficient transpeptidase with efficiency up to 1,340,000 M^−1^s^−1^, and to VyPAL2, OaAEP1, AtLegγ, and HaAEP1, which have intermediate to low transpeptidase efficiency. Enzymatic assays have shown that CeAEP1 favours hydrolysis over transpeptidation in a non-native substrate (Bernath-Levin et al., 2015). A few structural features of AEPs have been explored by several research groups, allowing us to better understand what makes CeAEP1 capable of transpeptidation, in addition to its proteolytic activity.

Residue 230 appears to be an important determinant for hydrolysis/transpeptidation. It belongs to a trio of residues (residue 229-331) that were designated Ligase-Activity Determinant 1 by Hemu et al. (2019) (**Supplemental Figure 10**). CeAEP1, like HaAEP1 and AtLegγ, is an inefficient transpeptidase with a glycine in this position. Transpeptidation efficiency was successfully enhanced by a cysteine to alanine mutation in this position in OaAEP1 (Yang et al., 2017). In contrast, a cysteine to valine/isoleucine mutation at the equivalent position in OaAEP1 abolished transpeptidase activity even though efficient transpeptidases butelase1 and VyPAL2 have valine and isoleucine respectively at this position (Yang et al., 2017). The effect of residue 230 on catalytic activity therefore appears to be dependent on a complex interplay with nearby residues. A glycine in this position, however, seems to be a good indicator of predominant protease activity. The same conclusion can be reached based on sequence comparison of other biochemically characterised AEPs (Hemu et al., 2019).

### Hydrophobic S2′ pocket facilitates transpeptidation

The hydrophobic nature of both the P2′ residue of pro-conA (isoleucine) and the S2′ pocket of CeAEP1 (Val163, Phe171, Tyr173) (**Figure 6B**) suggests that CeAEP1 might bind to the peptide residues C-terminal of the cleavage site. This would be crucial for preventing water molecules from entering the active site, therefore facilitating aminolysis instead of hydrolysis (Bernath-Levin et al., 2015; Zauner et al., 2018a). Consistent with this hypothesis, hydrophobic residues are present in the S2′ site of HaAEP1 (Val183, Val193), AtLegγ (Val182, Tyr190, Tyr192), OaAEP1 (Val180, Tyr188, Tyr190), VyPAL2 (Tyr185, Tyr187) and butelase1 (Val170, Tyr178, Ala180), and at the P2′ residue of their native substrate, SFTI-1 in sunflower (P2′ leucine), kalata-B1 in *O. affinis* (P2′ leucine), *V. yedoensis* cyclotide precursors (P2′ leucine) and *C. ternatea* cyclotide precursors (P2′ valine). There is no known peptide/protein in *A. thaliana* that undergoes AEP-mediated transpeptidation, but AtLegγ has also been shown to perform transpeptidation on non-native substrates with valine as the P2′ residue (Zauner et al., 2018b). Molecular dynamic simulations performed on VyPAL2 also showed that the S2′ pocket favours hydrophobic P2′ residues (Hemu et al., 2019). Gly165 conservation in the S2′ site appears to be essential to maintain a cavity in this hydrophobic S2′ pocket (**Figure 6B; Supplemental Figure 10**) as a glycine to serine mutation in HaAEP1 was shown to severely affect catalytic activity (Haywood et al., 2018).

### Facilitating N-terminal nucleophilic attack for transpeptidase activity

Previously, we proposed that residues around the catalytic cysteine facilitate N-terminal nucleophilic attack in transpeptidation (Haywood et al., 2018). This includes Glu201, Pro161, Gln225 and Asn227 of CeAEP1 (**Figure 6C**). Although *in silico* docking studies show that Glu201 forms an ionic bond with the incoming N-terminal nucleophile (Zauner et al., 2018a), this residue does not appear to be a major determinant of transpeptidation efficiency as glutamate is found in this position in AEPs that have differing transpeptidase efficiencies (**Supplemental Figure 10**). The nearby Gly160 (**Figure 6C**), which is one of the two residues designated as Ligase-Activity Determinant 2 by Hemu et al. (2019), is a highly conserved residue as most plant AEPs have a Gly or Ala at this residue (**Supplemental Figure 10**). At this position, a tyrosine to glycine mutation in VyPAL3 and a tyrosine to alanine mutation in *Viola Canadensis* AEP were shown to enhance the catalytic activity of transpeptidation. Any side chain bigger than alanine at residue 160 may therefore sterically interfere with the binding of peptide substrate residues C-terminal of the cleavage site, allowing water to enter to complete the cleavage reaction by hydrolysis.

Pro161, Gln225 and Asn227, on the other hand, have been hypothesised to sterically hinder the incoming N-terminus from interacting with Glu201 to initiate transpeptidation (Haywood et al., 2018). These three residues lining the active site are the same in CeAEP1 and HaAEP1, both of which are inefficient transpeptidases. In contrast, the smaller side chains in butelase1 (Gly167, Ala168, Gly232, Ser234) likely contribute to its high transpeptidase efficiency, and is also likely why it will accept most N-terminal amino acids for transpeptidation (Nguyen et al., 2014). The intermediate transpeptidation efficiency of AtLegγ, OaAEP1, and VyPAL2 relative to CeAEP1, HaAEP1 and butelase1 corroborates with the intermediate amino acid sizes (AtLegγ: Gly179, Pro180, Glu244, Ser246; OaAEP1: Ala177, Ala178, Thr242, Ser246; VyPAL2: Ala174, Pro175, Thr239, Gly241).

Although Gln225 and Asn227 in CeAEP1 and HaAEP1 may sterically interfere with transpeptidation, their long, polar side chains may allow for redundancy in interacting with the N-terminal nucleophile, as was observed in the absence of Glu201 (Haywood et al., 2018).

## Conclusion

Our investigations show that the mature form of conA is more stable than its precursor, pro-conA, revealing a functional consequence for the only known naturally-occurring circular permutation, which was discovered about 30 years ago. Structural evidence shows that no change occurs at the carbohydrate binding site, and ITC analysis shows that monosaccharide binding is not affected. The difference in crystal packing between pro-conA and conA indicates that changes induced by circular permutation occur at the quaternary rather than the tertiary level. Using purified recombinant proteins, we show that jack bean AEP is capable of carrying out both cleavage and transpeptidation reactions to circularly permutate pro-conA. Although CeAEP1 has been shown to be predominantly a protease, this was determined with a non-native substrate (Bernath-Levin et al., 2015). Here, structural analysis and recently-generated knowledge from other AEP structures allow us to understand how a protease like CeAEP1 is capable of transpeptidation. Although CeAEP1 is capable of conA circular permutation, cDNA analysis suggests the presence of AEP isoenzymes with unknown relative abundance (Takeda et al., 1994), so there is possible redundancy or involvement of another dedicated protease/transpeptidase for conA circular permutation *in vivo*.

**Table 1.**
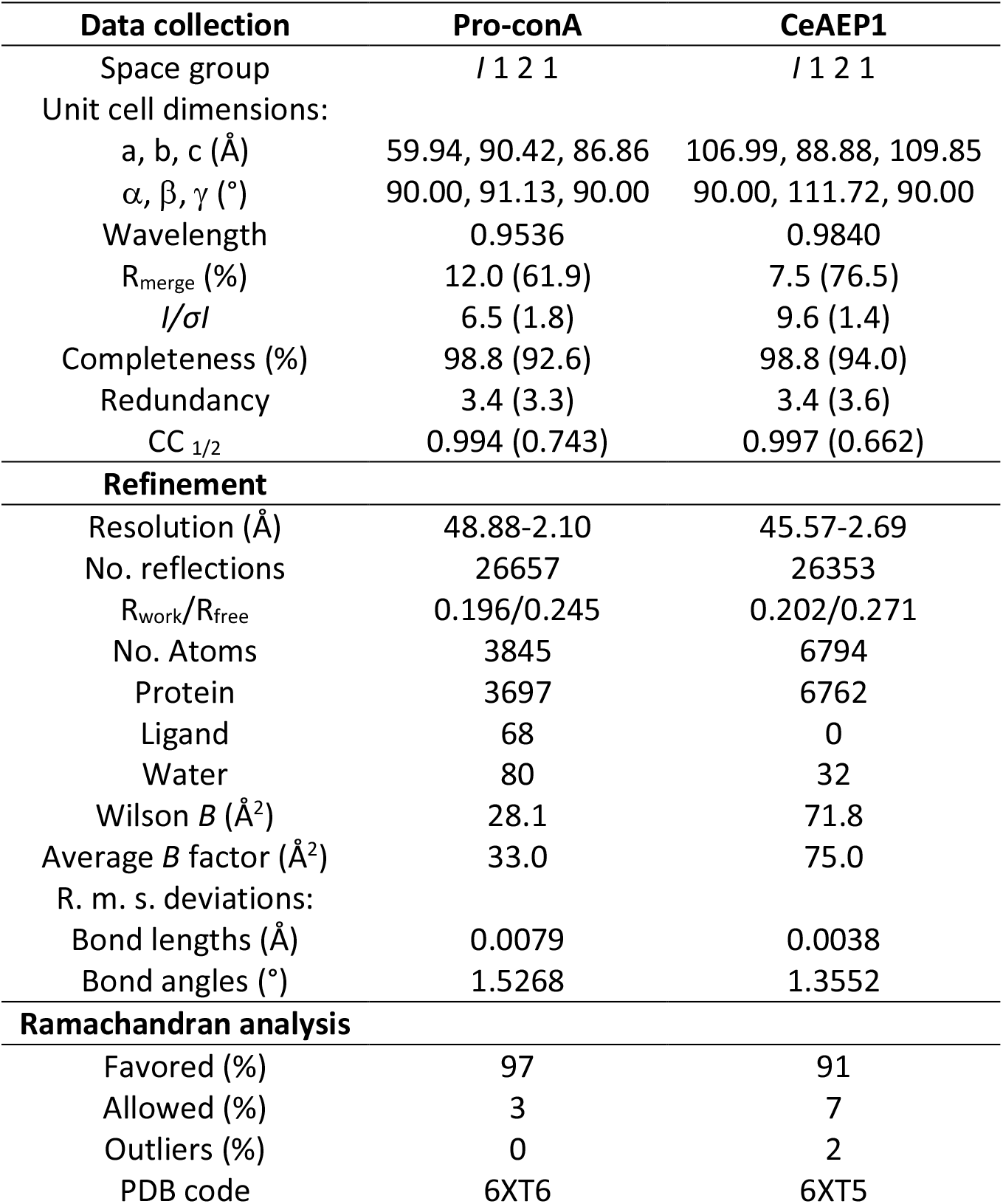
Summary of crystallographic data and refinement statistics for pro-conA and CeAEP1. Values in parentheses are for the highest resolution shell.

## Acknowledgements

This work was supported in part by Australian Research Council (ARC) grant DP160100107 to J.S.M. This research was undertaken in part using the MX2 beamline at the Australian Synchrotron, part of ANSTO, and made use of the Australian Cancer Research Foundation (ACRF) detector. J.H. was supported by an ARC Discovery Early Career Researcher Award DE180101445. S.G.N. was supported by the Australian Research Training Program.

## Supplemental Information

### Supplemental Figures

**Supplemental Figure 1.**
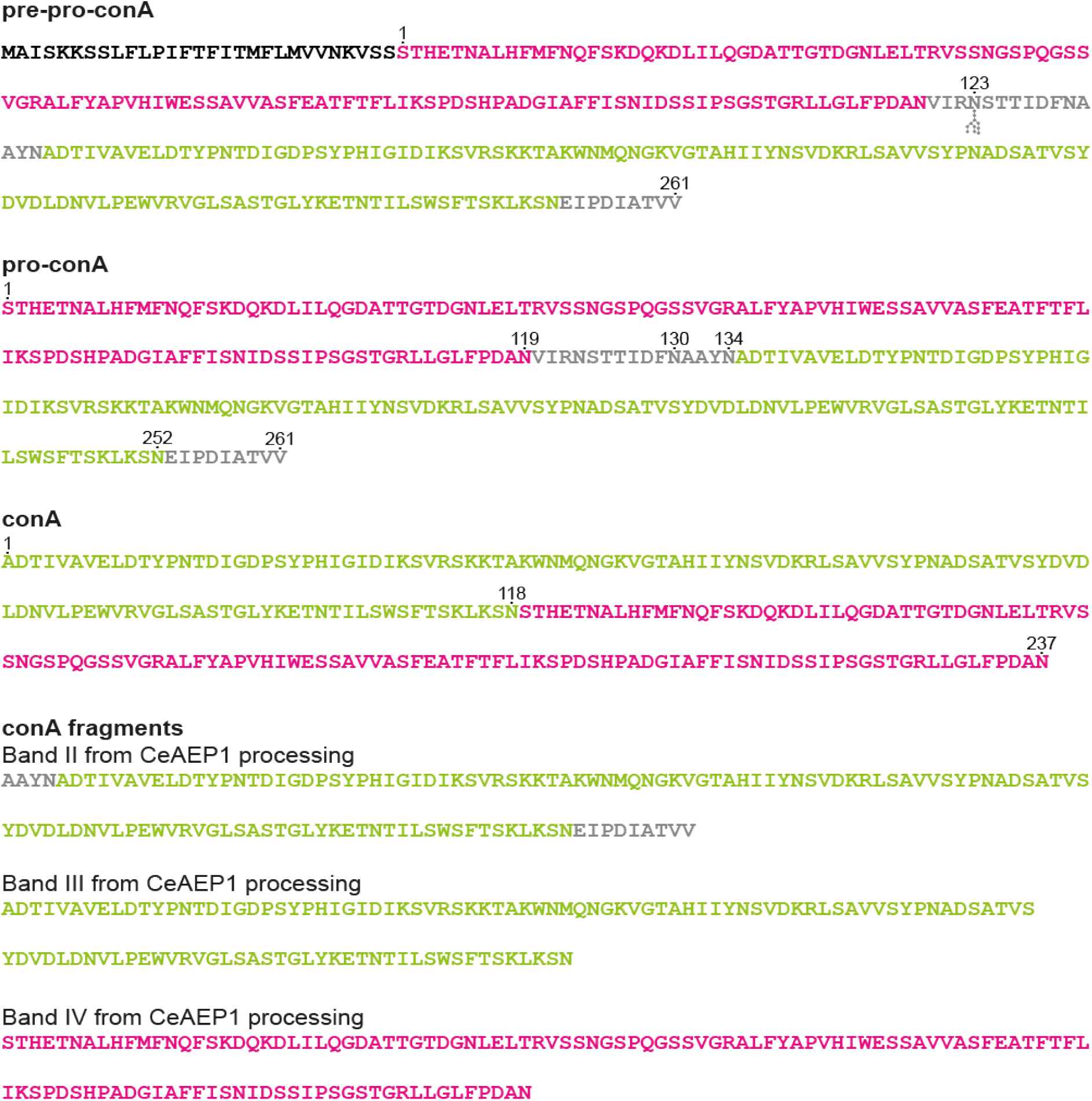
Sequence from pre-pro-conA to conA and conA fragments. Pre-pro-conA where chain B (pink) forms the N-terminal half and chain A (green) forms the C-terminal half of the protein. Pre-pro-conA contains an N-linked glycan group (grey ball-and-stick) at N123. ER-signal (black) is cleaved off to form pro-conA. N-linked glycan is typically included when describing pro-conA. For brevity, we describe deglycosylated conA as just conA. Residues cleaved off during circular permutation to form conA are in grey. Residue numbering for pre-pro-conA starts after N-terminal ER-signal to match pro-conA numbering. Residues renumbered in conA as chain A is circularly permuted to become the N-terminal half and chain B becomes the C-terminal half. Other than full-length conA, CeAEP1 processing *in vitro* produces conA fragments, labelled band II-IV on SDS-PAGE analysis (**Fig. 3**), which correlates to those observed in jack bean seeds (Sheldon et al., 1996).

**Supplemental Figure 2.**
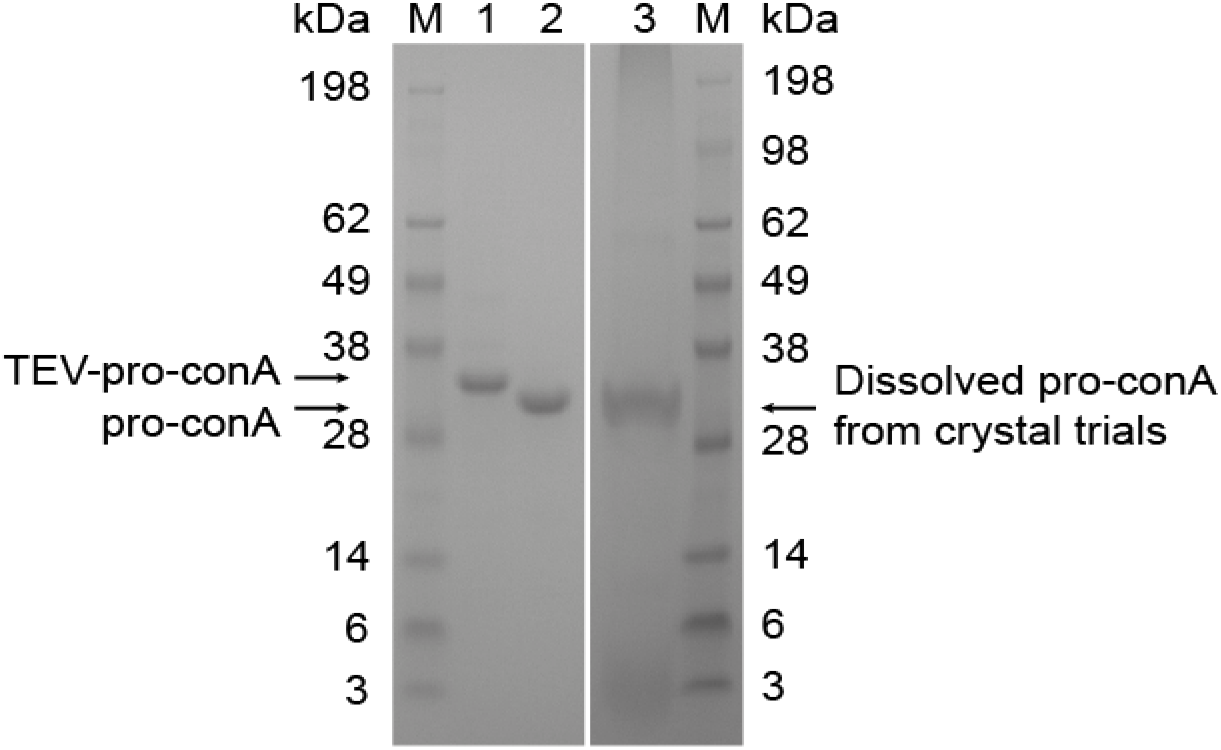
Recombinant pro-conA purification and confirming full-length protein after crystal trials. His-tagged TEV-pro-conA was purified on a nickel column (Lane 1) and cleaved by TEV protease to remove the N-terminal His-tag, reconstituting the native N-terminus in recombinant pro-conA. Recombinant pro-conA was purified from TEV protease using reverse His-tag purification on nickel column (Lane 2). SDS-PAGE confirming pro-conA is a full-length monomer despite weak electron density in the 15-amino acid intervening peptide that could have been interpreted as spurious cleavage by other enzymes. Recombinant pro-conA from crystal trials was dissolved in MOPS storage buffer, run on an SDS-PAGE gel and Coomassie stained.

**Supplemental Figure 3.**
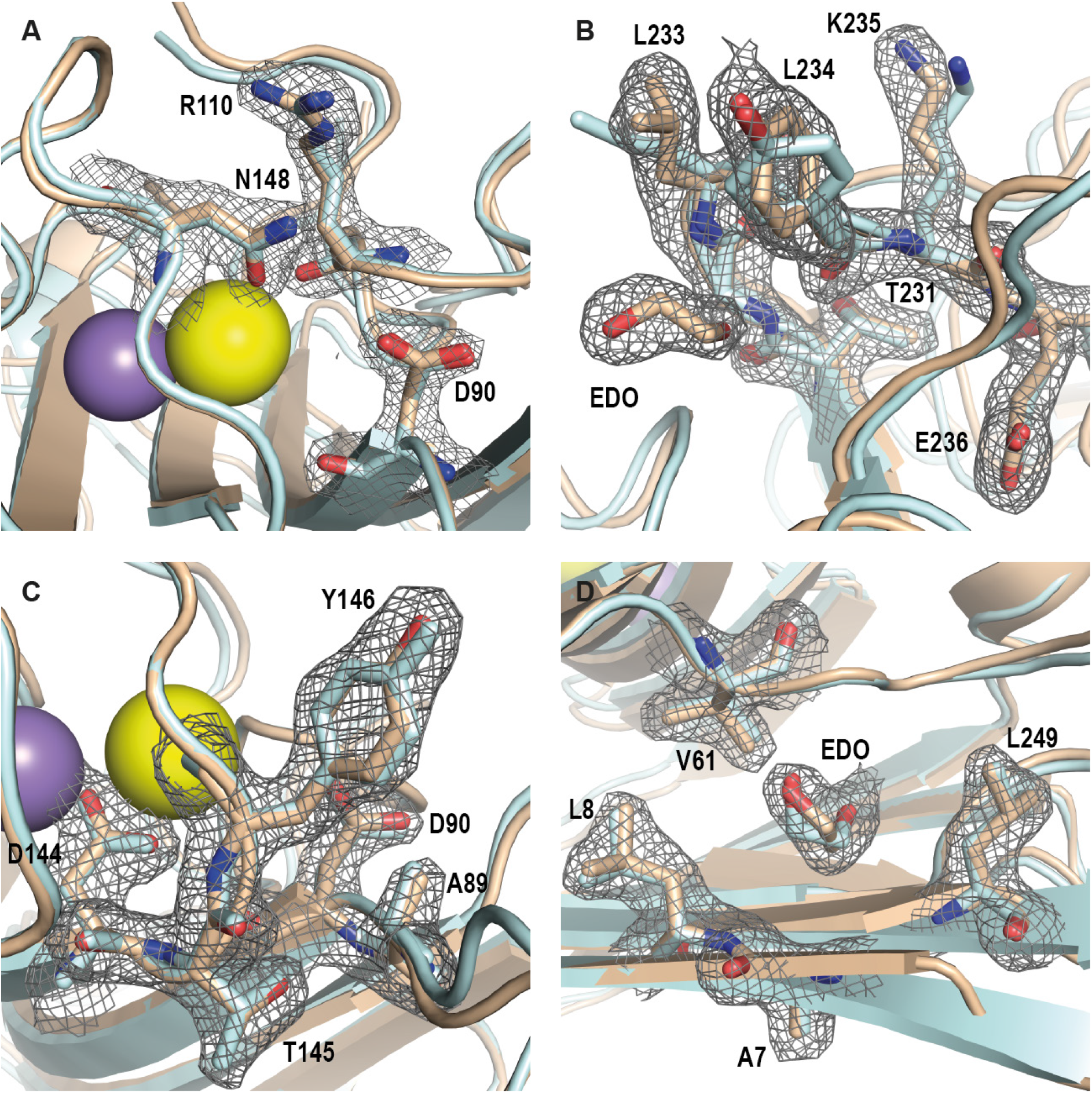
conA functional sites remain unchanged after circular permutation. (**A**) Monosaccharide binding site: Asp90, Arg110 and Asn148 in pro-conA; Asp208, Arg228 and Asn14 in conA (PDB: 1JBC). (**B**) Monosaccharide specificity loop: Thr231-Glu236 in pro-conA; Thr97-Glu102 in conA (PDB: 1JBC). (**C**) Key residues in locked/active state of pro-conA and conA: Ala89, Asp90, Asp144, Thr145, Tyr146 in pro-conA; Ala207, Asp208, Asp10, Thr11, Tyr12 in conA (PDB:1JBC). (**D**) Hypothesised hydrophobic cavity: Ala7, Leu8, Val61 and Leu249 in pro-conA; Ala125, Leu126, Val 179 and Leu115 in conA (PDB: 4PF5). Pro-conA (wheat); conA (pale cyan); calcium (yellow sphere); manganese (purple); ethane diol (EDO). Electron density maps (2*F*_obs_-*F*_calc_) for key amino acid residues are contoured at the 1σ level. PDB: 4PF5 used for comparing hydrophobic cavity in panel **D** as it was similarly crystallised with EDO in the cavity. The relative positions of these functional sites are highlighted in **Figure 2a.**

**Supplemental Figure 4.**
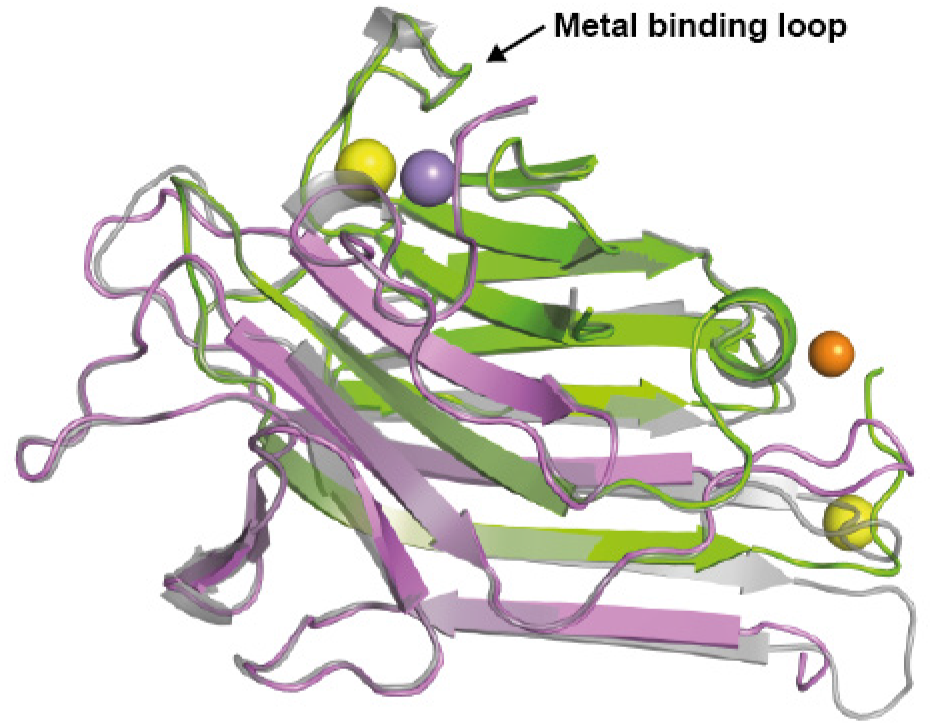
Pro-conA and conA have similar structures. Pro-conA and conA (PDB: 1JBC) have an r. m. s. d. of 1.2 Å. Overlay of pro-conA (pink and green as described in **Fig. 2**) and conA (grey). Calcium (yellow spheres) and manganese (purple sphere) ions in the ‘metal binding loop’ are perfectly overlaid between pro-conA and conA. Calcium and zinc (orange sphere) outside of the ‘metal binding loop’ are only present in pro-conA due to crystal conditions.

**Supplemental Figure 5.**
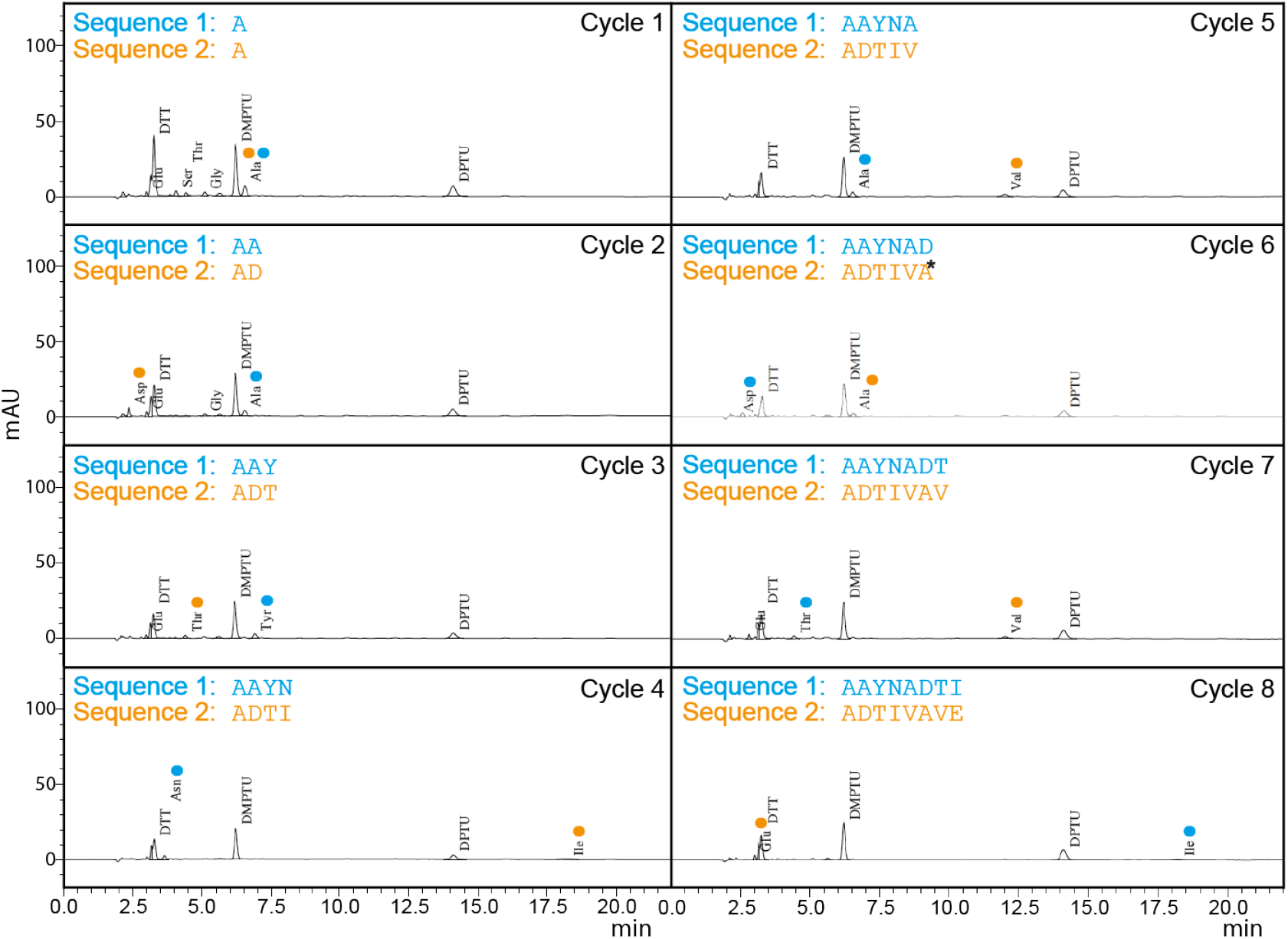
N-terminal Edman degradation of pro-conA processed by recombinant CeAEP1. N-terminal Edman degradation was performed on the protein band highlighted in **Fig. 3A**. A search for all possible permutations from Edman degradation indicated presence of two overlapping sequences (text in blue and orange, respectively) that are present in the conA sequence and underlined in **Fig. 3B**. DMPTU (dimethylphenylthiourea) and DPTU (diphenylthiourea) are normal by-products of the Edman degradation process. *Confidence for Ala in Cycle 6 is low because absorbance for Ala in Cycle 6 is lower than in Cycle 5, and so could be due to carry over from Cycle 5.

**Supplemental Figure 6.**
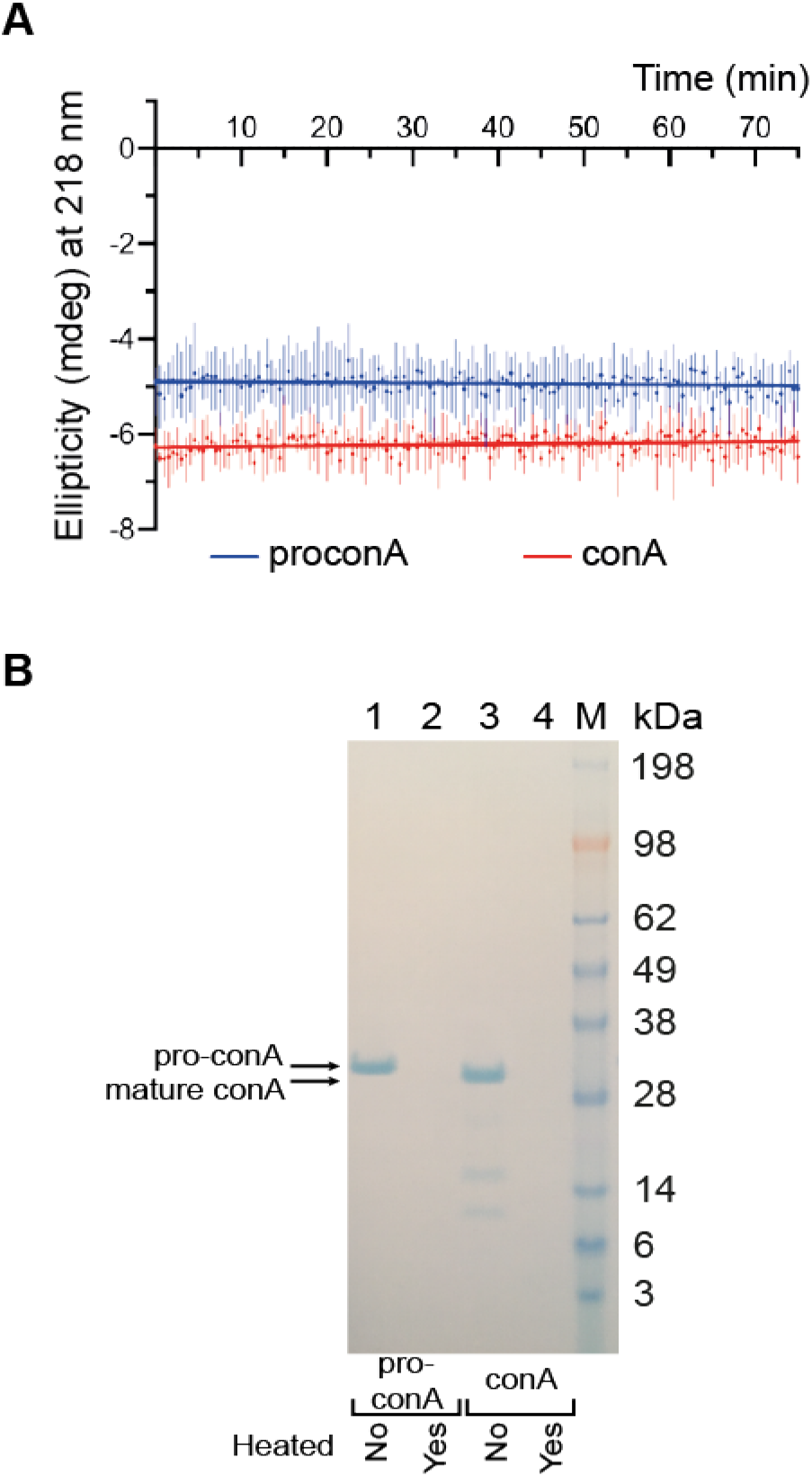
Pro-conA and conA stability with and without heat treatment. (**A**) No-heat controls show no dramatic changes to ellipticity at 218 nm for pro-conA and conA when maintained at 25 °C for the duration of the heat analysis (75 minutes) in **Fig. 4A**, indicating that pro-conA and conA are stable without heating. (**B**) SDS-PAGE analysis of pro-conA and conA after heat analysis by CD (Lanes 2 and 4) and after no-heat control analysis (Lanes 1 and 3). To determine if any soluble proteins remained, samples were centrifuged at 15,000 × g for 10 minutes and supernatant analysed on SDS-PAGE. Pro-conA (Lane 2) and conA (Lane 4). Pro-conA (Lane 1) and conA (Lane 3) no-heat controls.

**Supplemental Figure 7.**
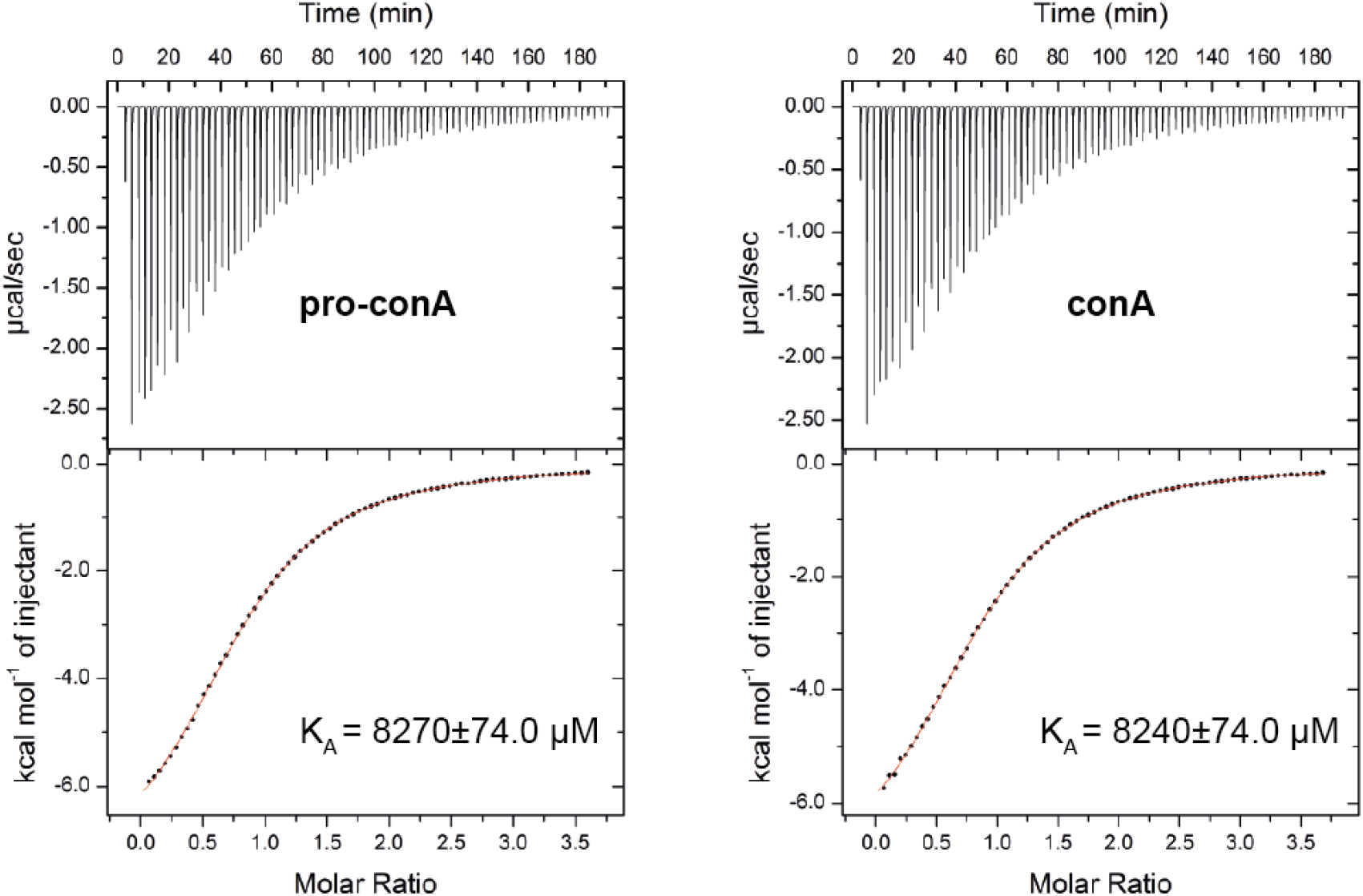
Circular permutation has no effect on conA binding to methyl-α-D-mannose. ITC analysis shows that pro-conA and conA have similar K_A_ (association constant). Pro-conA and conA were titrated with methyl-α-D-mannose (injectant) at 25 °C. Top panels display raw data for 76 0.5 μL-injections. Data points for the integrated curves are displayed in the bottom panel. Baseline and integration ranges were determined automatically by the Origin® software (version 2002, OriginLab Corporation) supplied by MicroCal.

**Supplemental Figure 8.**
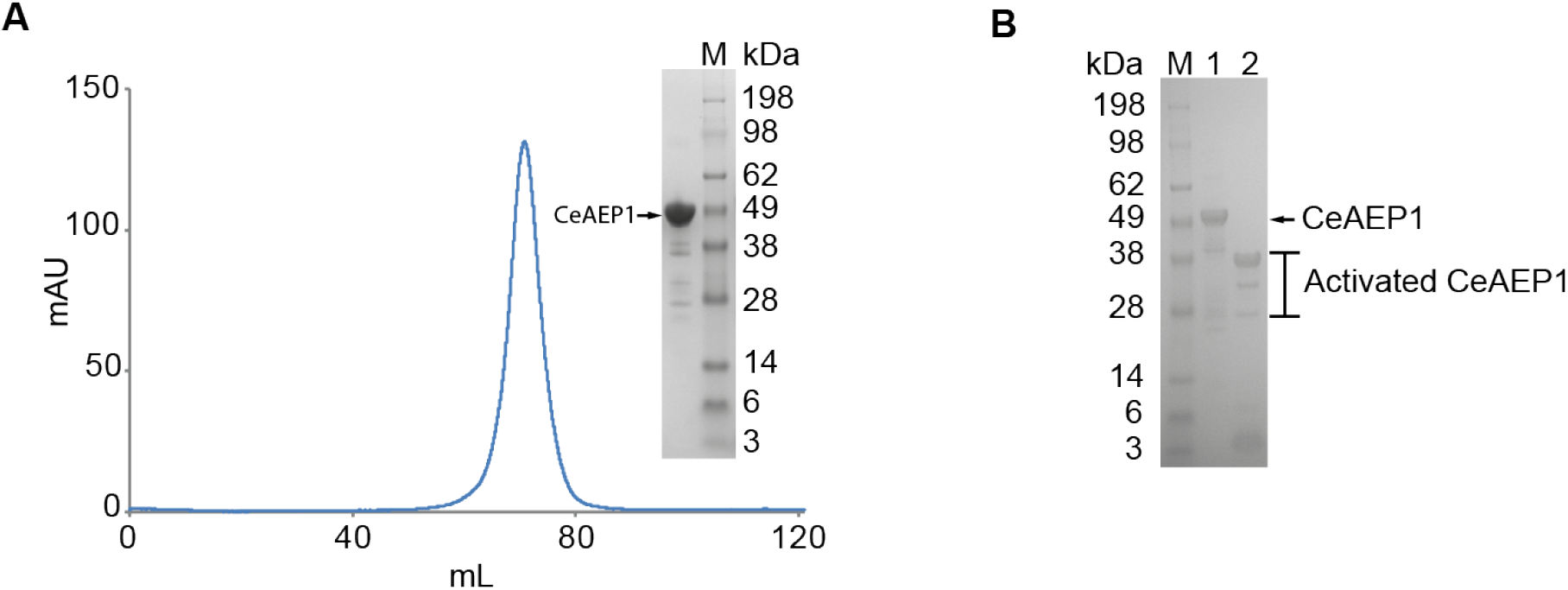
Recombinant CeAEP1 purification and activation. (**A**) Size exclusion chromatograph of CeAEP1 using HiLoad 16/600 Superdex 200. Inset: SDS-PAGE analysis of collected peak fraction. Lower MW proteins present after size exclusion possibly due to autocatalytic cleavage occurring at high protein concentration. (**B**) CeAEP1 activated by autocatalytic cleavage at pH 4. Lane 1: Nickel-purified CeAEP1. Lane 2: CeAEP1 dialysed at pH 4 for 4 hours then dialysed to pH 6.5. Multiple bands indicate that autocatalytic cleavage occurs at multiple sites, which is typical for AEPs (Yang et al., 2017; Haywood et al., 2018; Hemu et al., 2019; James et al., 2019).

**Supplemental Figure 9.**
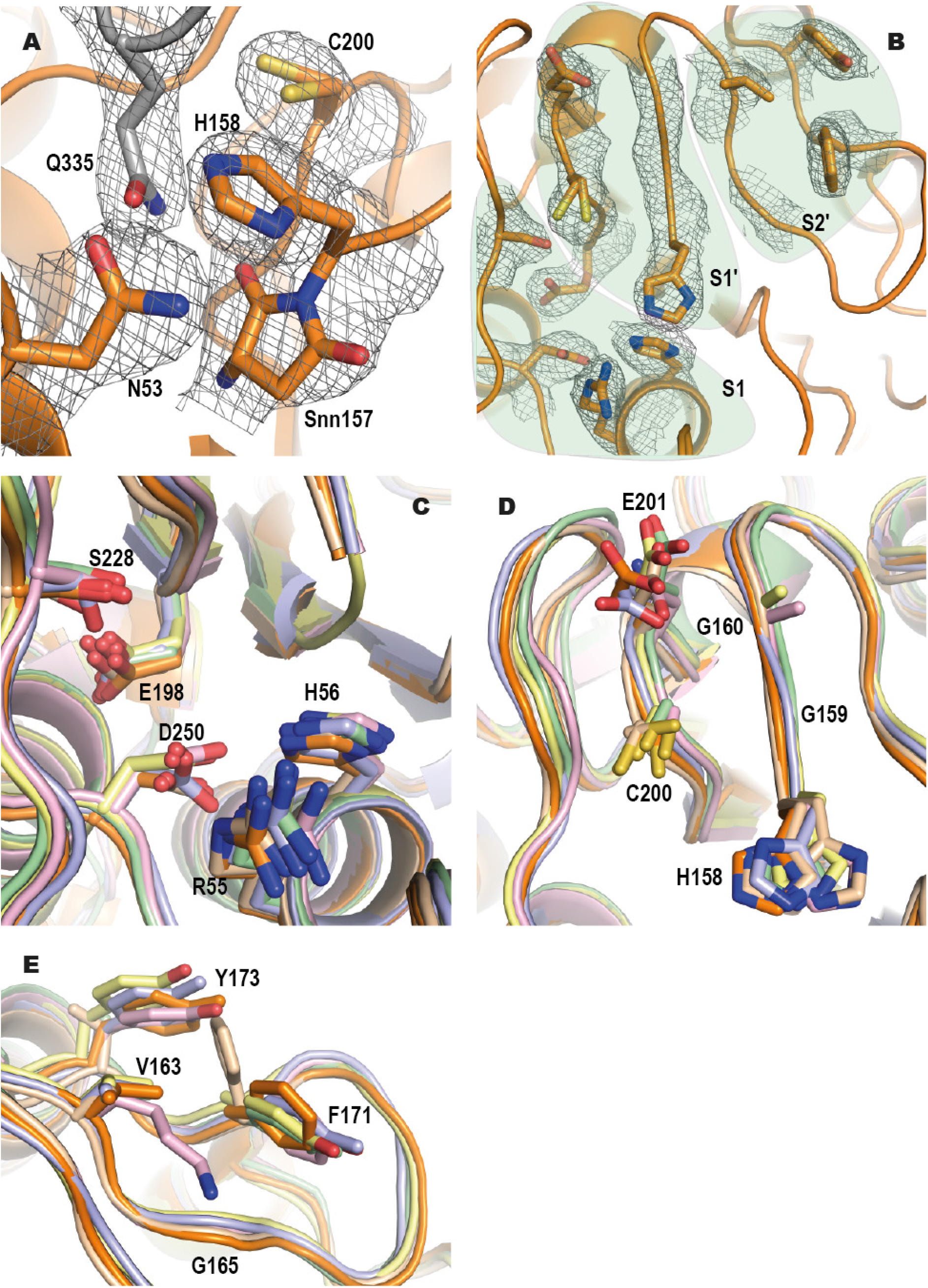
CeAEP1 residues involved in catalytic activity. Electron density maps (2 F_obs_ – F_calc_) contoured at 1σ level (**A**) Catalytic triad (N53, H158, C200) and succinimide intermediate (Snn157) displayed as orange sticks; glutamine residue from cap domain (Q335) displayed as grey sticks. (**B**) Substrate binding pockets viewed from above the core domain with residues in S1, S1′ and S2′ pockets labelled and shaded accordingly. Structural alignment of S1 pocket (**C**), S1′ pocket (**D**) and S2′ pocket (**E**). CeAEP1 (orange), HaAEP1 (wheat), AtLEGγ (light blue), OaAEP1 (pale yellow), VyPAL2 (light pink), butelase1 (pale green). CeAEP1 residues labelled. Catalytic cysteine of AtLEGγ (C200 in CeAEP1) not displayed as it was crystallised as methylated cysteine.

**Supplemental Figure 10.**
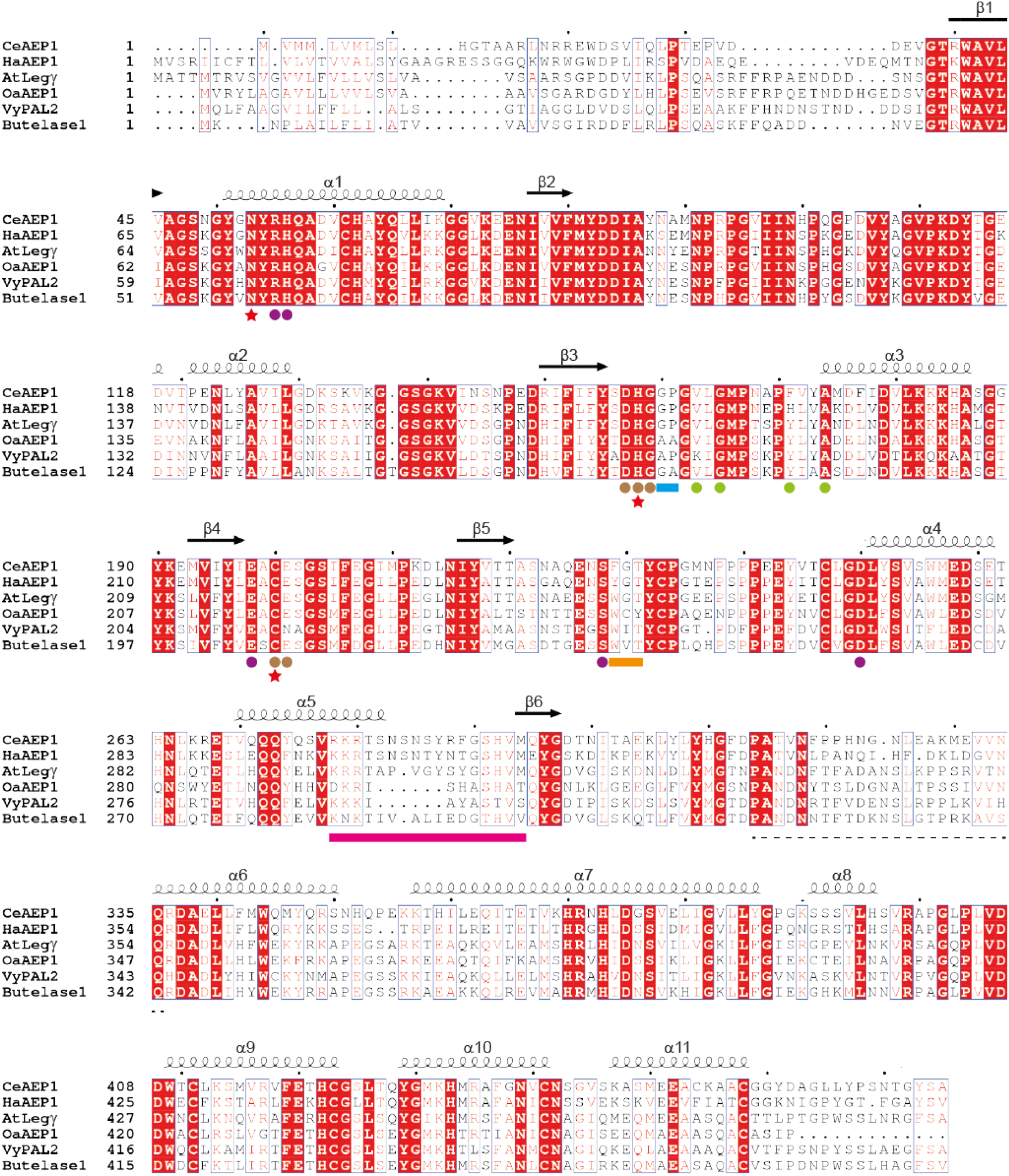
AEP Alignment. Protein sequence alignment of CeAEP1 (Uniprot: P49046), HaAEP1 (Uniprot: A0A0G2RI59), AtLegγ (Q39119), OaAEP1 (Uniprot: A0A0N9JZ32), VyPAL2 (Uniprot: 0A509GV09), and butelase1 (Uniprot: A0A060D9Z7). CeAEP1 major secondary structure is labelled and numbered above each row. Catalytic triad (red stars); S1 pocket (purple circles); S1′ pocket (brown circles); S2′ pocket (green circles); regions so-called Ligase-Activity Determinant 1 (orange bar); Ligase-Activity Determinant 2 (blue bar); Marker of Ligase Activity (pink bar). Flexible linker region between ‘core’ and ‘cap’ domains (dash line).

### Supplemental Tables

**Supplemental Table 1:** Interactions analysis using PDBePISA.

| Interactions | Buried surface area (Å<sup>2</sup>) | No. of interface residues | No. of salt bridges | No. of hydrogen bonds |
| --- | --- | --- | --- | --- |
| <sup>a</sup>pro-conA – subunits I and III | 733 | 24 | 5 | 6 |
| <sup>a</sup>conA – subunits I and III | 1034 | 31 | 8 | 12 |
| <sup>b</sup>pro-conA – subunits I and II | 1012 | 30 | 0 | 16 |
| <sup>b</sup>conA – subunits I and II | 1124 | 31 | 0 | 24 |

<sup>a</sup>Interactions between subunits I and III in a tetrameric complex (See **Figure2B**).
<sup>b</sup>Interactions between subunits I and II in a tetrameric complex (See **Figure2B**).

**Supplemental Table 2:**
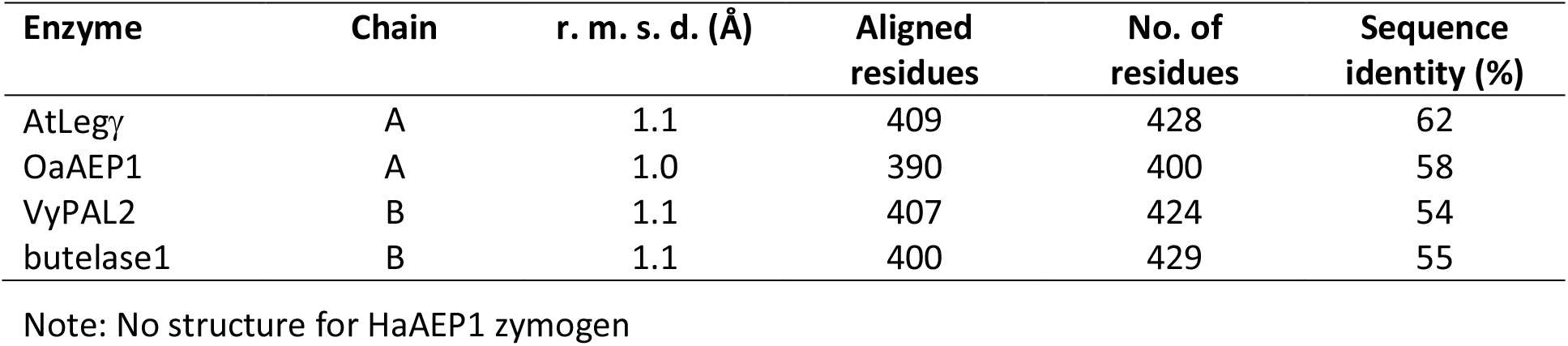
r. m. s. d. of Cα-atoms of CeAEP1 zymogen and its homologs.

**Supplemental Table 3:**
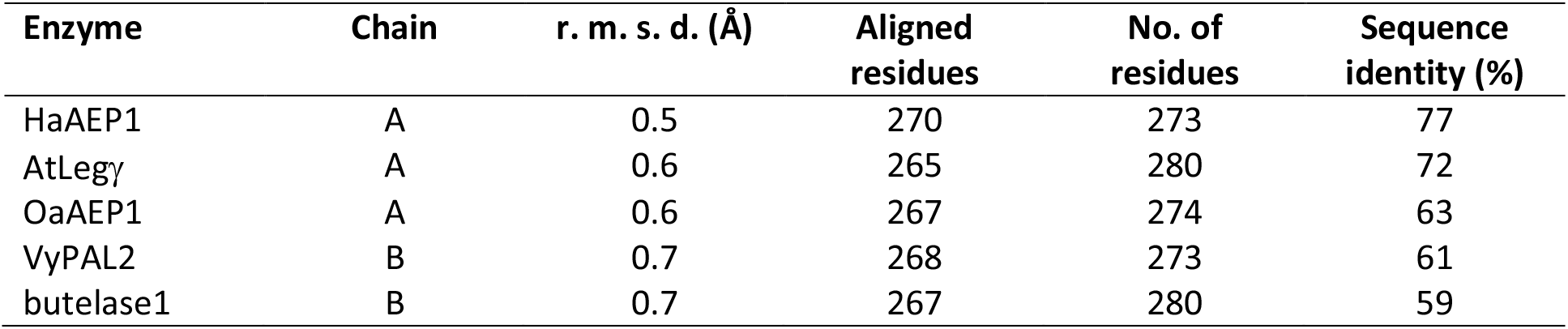
r. m. s. d. of Cα-atoms of CeAEP1 core domain and its homologs.

## References

Abe, Y., Shirane, K., Yokosawa, H., Matsushita, H., Mitta, M., Kato, I., and Ishii, S. (1993). Asparaginyl endopeptidase of jack bean seeds. Purification, characterization, and high utility in protein sequence analysis. J. Biol. Chem. 268, 3525–3529.

Adachi, M., Takenaka, Y., Gidamis, A.B., Mikami, B., and Utsumi, S. (2001). Crystal structure of soybean proglycinin A1aB1b homotrimer. J. Mol. Biol. 305, 291–305.

Adachi, M., Kanamori, J., Masuda, T., Yagasaki, K., Kitamura, K., Mikami, B., and Utsumi, S. (2003). Crystal structure of soybean 11S globulin: glycinin A3B4 homohexamer. Proc. Natl. Acad. Sci. USA 100, 7395–7400.

Bernath-Levin, K., Nelson, C., Elliott, A.G., Jayasena, A.S., Millar, A.H., Craik, D.J., and Mylne, J.S. (2015). Peptide macrocyclization by a bifunctional endoprotease. Chem. Biol. 22, 571–582.

Bernhard, W., and Avrameas, S. (1971). Ultrastructural visualization of cellular carbohydrate components by means of concanavalin A. Exp. Cell Res. 64, 232–236.

Bezerra, E.H., Rocha, B.A., Nagano, C.S., Bezerra Gde, A., Moura, T.R., Bezerra, M.J., Benevides, R.G., Sampaio, A.H., Assreuy, A.M., Delatorre, P., and Cavada, B.S. (2011). Structural analysis of ConBr reveals molecular correlation between the carbohydrate recognition domain and endothelial NO synthase activation. Biochem. Biophys. Res. Commun. 408, 566–570.

Bliven, S., and Prlić, A. (2012). Circular permutation in proteins. PLoS Comp. Biol. 8, e1002445.

Blumberg, S., and Tal, N. (1976). Effect of divalent metal ions on the digestibility of concanavalin a by endopeptidases. Biochim. Biophys. Acta 453, 357–364.

Bouckaert, J., Dewallef, Y., Poortmans, F., Wyns, L., and Loris, R. (2000). The structural features of concanavalin A governing non-proline peptide isomerization. J. Biol. Chem. 275, 19778–19787.

Bowles, D.J., and Pappin, D.J. (1988). Traffic and assembly of concanavalin A. Trends Biochem. Sci. 13, 60–64.

Bowles, D.J., Marcus, S.E., Pappin, D.J., Findlay, J.B., Eliopoulos, E., Maycox, P.R., and Burgess, J. (1986). Posttranslational processing of concanavalin A precursors in jackbean cotyledons. J. Cell Biol. 102, 1284–1297.

Brewer, C.F., Brown, R.D., 3rd, and Koenig, S.H. (1983). Metal ion binding and conformational transitions in concanavalin A: a structure-function study. J. Biomol. Struct. Dyn. 1, 961–997.

Brown, R.D., 3rd, Brewer, C.F., and Koenig, S.H. (1977). Conformation states of concanavalin A: kinetics of transitions induced by interaction with Mn2+ and Ca2+ ions. Biochemistry 16, 3883–3896.

Carrington, D.M., Auffret, A., and Hanke, D.E. (1985). Polypeptide ligation occurs during post-translational modification of concanavalin A. Nature 313, 64–67.

Cavada, B.S., Pinto-Junior, V.R., Osterne, V.J.S., and Nascimento, K.S. (2018). ConA-like lectins: High similarity proteins as models to study structure/biological activities relationships. Int. J. Mol. Sci. 20.

Chervenak, M.C., and Toone, E.J. (1995). Calorimetric Analysis of the Binding of Lectins with Overlapping Carbohydrate-Binding Ligand Specificities (Vol 34, Pg 5685, 1995). Biochemistry 34, 7966–7966.

Cunningham, B.A., Hemperly, J.J., Hopp, T.P., and Edelman, G.M. (1979). Favin versus concanavalin A: Circularly permuted amino acid sequences. Proc. Natl. Acad. Sci. USA 76, 3218–3222.

Delatorre, P., Rocha, B.A., Souza, E.P., Oliveira, T.M., Bezerra, G.A., Moreno, F.B., Freitas, B.T., Santi-Gadelha, T., Sampaio, A.H., Azevedo, W.F., Jr., and Cavada, B.S. (2007). Structure of a lectin from *Canavalia gladiata* seeds: new structural insights for old molecules. BMC Struct. Biol. 7, 52.

Doyle, R.J., Thomasson, D.L., and Nicholson, S.K. (1976). Stabilization of concanavalin A by metal ligands. Carbohydr Res 46, 111–118.

Du, J., Yap, K., Chan, L.Y., Rehm, F.B.H., Looi, F.Y., Poth, A.G., Gilding, E.K., Kaas, Q., Durek, T., and Craik, D.J. (2020). A bifunctional asparaginyl endopeptidase efficiently catalyzes both cleavage and cyclization of cyclic trypsin inhibitors. Nat. Commun. 11, 1575.

Dwyer, J.M., and Johnson, C. (1981). The use of concanavalin A to study the immunoregulation of human T cells. Clin. Exp. Immunol. 46, 237–249.

Edelman, G.M., and Wang, J.L. (1978). Binding and functional properties of concanavalin A and its derivatives. III. Interactions with indoleacetic acid and other hydrophobic ligands. J. Biol. Chem. 253, 3016–3022.

Einhoff, W., Fleischmann, G., Freier, T., Kummer, H., and Rüdiger, H. (1986). Interactions between lectins and other components of leguminous protein bodies. Biol. Chem. Hoppe-Seyler 367, 15–25.

Emmerich, C., Helliwell, J.R., Redshaw, M., Naismith, J.H., Harrop, S.J., Raftery, J., Kalb, A.J., Yariv, J., Dauter, Z., and Wilson, K.S. (1994). High-resolution structures of single-metal-substituted concanavalin A: the Co,Ca-protein at 1.6 A and the Ni,Ca-protein at 2.0 A. Acta Crystallogr. D 50, 749–756.

Emsley, P., Lohkamp, B., Scott, W.G., and Cowtan, K. (2010). Features and development of Coot. Acta Crystallogr. D 66, 486–501.

Faye, L., and Chrispeels, M.J. (1987). Transport and Processing of the Glycosylated Precursor of Concanavalin-a in Jack-Bean. Planta 170, 217–224.

François-Heude, M., Méndez-Ardoy, A., Cendret, V., Lafite, P., Daniellou, R., Ortiz Mellet, C., García Fernández, J.M., Moreau, V., and Djedaïni-Pilard, F. (2015). Synthesis of high-mannose oligosaccharide analogues through click chemistry: true functional mimics of their natural counterparts against lectins? Chemistry 21, 1978–1991.

Gebhard, L.G., Risso, V.A., Santos, J., Ferreyra, R.G., Noguera, M.E., and Ermácora, M.R. (2006). Mapping the distribution of conformational information throughout a protein sequence. J. Mol. Biol. 358, 280–288.

Goldenberg, D.P., and Creighton, T.E. (1983). Circular and circularly permuted forms of bovine pancreatic trypsin inhibitor. J. Mol. Biol. 165, 407–413.

Goldstein, J.I., Winter, C.H., and Poretz, D.R. (1997). Plant lectins: tools for the study of complex carbohydrates. In Glycoproteins II, J. Montreuil, J.F. G. Vliegenthart, and H. Schachter, eds (Elsevier B.V.), pp. 403–474.

Gunther, G.R., Wang, J.L., Yahara, I., Cunningham, B.A., and Edelman, G.M. (1973). Concanavalin A derivatives with altered biological activities. Proc. Natl. Acad. Sci. USA 70, 1012–1016.

Hamodrakas, S.J., Kanellopoulos, P.N., Pavlou, K., and Tucker, P.A. (1997). The crystal structure of the complex of concanavalin A with 4'-methylumbelliferyl-alpha-D-glucopyranoside. J. Struct. Biol. 118, 23–30.

Harris, K.S., Guarino, R.F., Dissanayake, R.S., Quimbar, P., McCorkelle, O.C., Poon, S., Kaas, Q., Durek, T., Gilding, E.K., Jackson, M.A., Craik, D.J., van der Weerden, N.L., Anders, R.F., and Anderson, M.A. (2019). A suite of kinetically superior AEP ligases can cyclise an intrinsically disordered protein. Sci. Rep. 9, 10820.

Haywood, J., Schmidberger, J.W., James, A.M., Nonis, S.G., Sukhoverkov, K.V., Elias, M., Bond, C.S., and Mylne, J.S. (2018). Structural basis of ribosomal peptide macrocyclization in plants. eLife 7, e32955.

Heinemann, U., and Hahn, M. (1995). Circular permutation of polypeptide chains: Implications for protein folding and stability. Prog. Biophys. Mol. Biol. 64, 121–143.

Hemu, X., Qiu, Y., Nguyen, G.K.T., and Tam, J.P. (2016). Total synthesis of circular bacteriocins by Butelase 1. J. Am. Chem. Soc. 138, 6968–6971.

Hemu, X., El Sahili, A., Hu, S., Wong, K., Chen, Y., Wong, Y.H., Zhang, X., Serra, A., Goh, B.C., Darwis, D.A., Chen, M.W., Sze, S.K., Liu, C.F., Lescar, J., and Tam, J.P. (2019). Structural determinants for peptide-bond formation by asparaginyl ligases. Proc. Natl. Acad. Sci. USA 116, 11737–11746.

Hendrix, R.W. (1991). Protein carpentry. Curr. Biol. 1, 71–73.

Herman, E.M., Shannon, L.M., and Chrispeels, M.J. (1985). Concanavalin A is synthesized as a glycoprotein precursor. Planta 165, 23–29.

Huet, M., and Claverie, J.M. (1978). Sedimentation Studies of Reversible Dimer-Tetramer Transition Kinetics of Concanavalin-A. Biochemistry 17, 236–241.

Jackson, M.A., Gilding, E.K., Shafee, T., Harris, K.S., Kaas, Q., Poon, S., Yap, K., Jia, H., Guarino, R., Chan, L.Y., Durek, T., Anderson, M.A., and Craik, D.J. (2018). Molecular basis for the production of cyclic peptides by plant asparaginyl endopeptidases. Nat. Commun. 9, 2411.

James, A.M., Haywood, J., Leroux, J., Ignasiak, K., Elliott, A.G., Schmidberger, J.W., Fisher, M.F., Nonis, S.G., Fenske, R., Bond, C.S., and Mylne, J.S. (2019). The macrocyclizing protease butelase 1 remains autocatalytic and reveals the structural basis for ligase activity. Plant J. 98, 988–999.

Jung, R., Scott, M.P., Nam, Y.-W., Beaman, T.W., Bassüner, R., Saalbach, I., Müntz, K., and Nielsen, N.C. (1998). The role of proteolysis in the processing and assembly of 11S seed globulins. The Plant Cell 10, 343–357.

Kabsch, W. (2010). Xds. Acta Crystallogr. D 66, 125–132.

Kalb, A.J., and Levitzki, A. (1968). Metal-binding sites of concanavalin A and their role in the binding of alpha-methyl d-glucopyranoside. Biochem. J. 109, 669–672.

Kanellopoulos, P.N., Tucker, P.A., Pavlou, K., Agianian, B., and Hamodrakas, S.J. (1996a). A triclinic crystal form of the lectin concanavalin A. J. Struct. Biol. 117, 16–23.

Kanellopoulos, P.N., Pavlou, K., Perrakis, A., Agianian, B., Vorgias, C.E., Mavrommatis, C., Soufi, M., Tucker, P.A., and Hamodrakas, S.J. (1996b). The crystal structure of the complexes of concanavalin A with 4’-nitrophenyl-alpha-D-mannopyranoside and 4’-nitrophenyl-alpha-D-glucopyranoside. J. Struct. Biol. 116, 345–355.

Krissinel, E., and Henrick, K. (2007). Inference of macromolecular assemblies from crystalline state. J. Mol. Biol. 372, 774–797.

Lagarda-Diaz, I., Guzman-Partida, A.M., and Vazquez-Moreno, L. (2017). Legume lectins: Proteins with diverse applications. Int. J. Mol. Sci. 18.

Laskowski, R.A., Jabłońska, J., Pravda, L., Vařeková, R.S., and Thornton, J.M. (2018). PDBsum: Structural summaries of PDB entries. Protein Sci. 27, 129–134.

Lis, H., and Sharon, N. (1998). Lectins: Carbohydrate-specific proteins that mediate cellular recognition. Chem. Rev. 98, 637–674.

Locke, A.K., Cummins, B.M., Abraham, A.A., and Cote, G.L. (2014). PEGylation of concanavalin A to improve its stability for an in vivo glucose sensing assay. Anal Chem 86, 9091–9097.

Loka, R.S., McConnell, M.S., and Nguyen, H.M. (2015). Studies of Highly-Ordered Heterodiantennary Mannose/Glucose-Functionalized Polymers and Concanavalin A Protein Interactions Using Isothermal Titration Calorimetry. Biomacromolecules 16, 4013–4021.

Loris, R., Hamelryck, T., Bouckaert, J., and Wyns, L. (1998). Legume lectin structure. Biochim. Biophys. Acta 1383, 9–36.

Macedo, M.L., das Gracas Machado Freire, M., da Silva, M.B., and Coelho, L.C. (2007). Insecticidal action of *Bauhinia monandra* leaf lectin (BmoLL) against *Anagasta kuehniella* (Lepidoptera: Pyralidae), *Zabrotes subfasciatus* and *Callosobruchus maculatus* (Coleoptera: Bruchidae). Comp. Biochem. Physiol. A Mol. Integr. Physiol. 146, 486–498.

Maeda, H., Hattori, H., and Kanoh, H. (1989). Conformational change and aggregation of concanavalin A at high temperatures. Int. J. Biol. Macromol. 11, 290–296.

McCubbin, W.D., Oikawa, K., and Kay, C.M. (1971). Circular dichroism studies on concanavalin A. Biochem. Biophys. Res. Commun. 43, 666–674.

McKenzie, G.H., Sawyer, W.H., and Nichol, L.W. (1972). The molecular weight and stability of concanavalin A. Biochim. Biophys. Acta 263, 283–293.

McPhillips, T.M., McPhillips, S.E., Chiu, H.-J., Cohen, A.E., Deacon, A.M., Ellis, P.J., Garman, E., Gonzalez, A., Sauter, N.K., Phizackerley, R.P., Soltis, S.M., and Kuhn, P. (2002). Blu-Ice and the Distributed Control System: software for data acquisition and instrument control at macromolecular crystallography beamlines. J. Synchrotron Radiat. 9, 401–406.

Meister, G.E., Kanwar, M., and Ostermeier, M. (2011). Circular permutation of proteins. In Protein Engineering Handbook, S. Lutz and U.T. Bornscheuer, eds (Wiley-VCH Verlag GmbH & Co.), pp. 453–471.

Melander, M., Åhman, I., Kamnert, I., and Strömdahl, A.C. (2003). Pea lectin expressed transgenically in oilseed rape reduces growth rate of pollen beetle larvae. Transgenic Res. 12, 555–567.

Min, W., and Jones, D.H. (1994). *In vitro* splicing of concanavalin A is catalyzed by asparaginyl endopeptidase. Nat. Struct. Mol. Biol. 1, 502–504.

Min, W., Dunn, A.J., and Jones, D.H. (1992). Non-glycosylated recombinant pro-concanavalin A is active without polypeptide cleavage. EMBO J. 11, 1303–1307.

Nguyen, G.K.T., Wang, S., Qiu, Y., Hemu, X., Lian, Y., and Tam, J.P. (2014). Butelase 1 is an Asx-specific ligase enabling peptide macrocyclization and synthesis. Nat. Chem. Biol. 10, 732–738.

Ogata, S., Muramatsu, T., and Kobata, A. (1975). Fractionation of glycopeptides by affinity column chromatography on concanavalin A-sepharose. J. Biochem. 78, 687–696.

Parkin, S., Rupp, B., and Hope, H. (1996). Atomic resolution structure of concanavalin A at 120 K. Acta Crystallogr. D 52, 1161–1168.

Peumans, W.J., and Van Damme, E.J. (1995). Lectins as plant defense proteins. Plant Physiol. 109, 347–352.

Ramis, C., Gomord, V., Lerouge, P., and Faye, L. (2001). Deglycosylation is necessary but not sufficient for activation of proconcanavalin A. J. Exp. Bot. 52, 911–917.

Saleemuddin, M., and Husain, Q. (1991). Concanavalin A: a useful ligand for glycoenzyme immobilization--a review. Enzyme Microb. Technol. 13, 290–295.

Schechter, I., and Berger, A. (1967). On the size of the active site in proteases. I. Papain. Biochem. Biophys. Res. Commun. 27, 157–162.

Senear, D.F., and Teller, D.C. (1981). Thermodynamics of concanavalin A dimer-tetramer self-association: sedimentation equilibrium studies. Biochemistry 20, 3076–3083.

Sharon, N., and Lis, H. (1990). Legume lectins--a large family of homologous proteins. FASEB J. 4, 3198–3208.

Sheldon, P.S., and Bowles, D.J. (1992). The glycoprotein precursor of concanavalin A is converted to an active lectin by deglycosylation. EMBO J. 11, 1297–1301.

Sheldon, P.S., Keen, J.N., and Bowles, D.J. (1996). Post-translational peptide bond formation during concanavalin A processing *in vitro*. Biochem. J. 320, 865–870.

Shoham, M., Kalb, A.J., and Pecht, I. (1973). Specificity of metal ion interaction with concanavalin A. Biochemistry 12, 1914–1917.

Smith, S.C., Johnson, S., Andrews, J., and McPherson, A. (1982). Biochemical characterization of canavalin, the major storage protein of jack bean. Plant Physiol. 70, 1199–1209.

Sumner, B.J. (1919). The globulins of the jack bean, *Canavalia ensiformis*. J. Biol. Chem. 37, 137–142.

Sumner, J.B., and Howell, S.F. (1936). Identification of Hemagglutinin of Jack Bean with Concanavalin A. J. Bacteriol. 32, 227–237.

Takeda, O., Miura, Y., Mitta, M., Matsushita, H., Kato, I., Abe, Y., Yokosawa, H., and Ishii, S.-I. (1994). Isolation and analysis of cDNA encoding a precursor of *Canavalia ensiformis* asparaginyl endopeptidase (legumain). J. Biochem. 116, 541–546.

Tang, T.M.S., Cardella, D., Lander, A.J., Li, X.F., Escudero, J.S., Tsai, Y.H., and Luk, L.Y.P. (2020). Use of an asparaginyl endopeptidase for chemoenzymatic peptide and protein labeling. Chem. Sci. 11, 5881–5888.

Topell, S., Hennecke, J., and Glockshuber, R. (1999). Circularly permuted variants of the green fluorescent protein. FEBS Lett. 457, 283–289.

Towbin, H., Staehelin, T., and Gordon, J. (1979). Electrophoretic transfer of proteins from polyacrylamide gels to nitrocellulose sheets: procedure and some applications. Proc. Natl. Acad. Sci. USA 76, 4350–4354.

Wang, J.L., Cunningham, B.A., and Edelman, G.M. (1971). Unusual fragments in the subunit structure of concanavalin A. Proc. Natl. Acad. Sci. USA 68, 1130–1134.

Winn, M. D., Ballard, C.C., Cowtan, K.D., Dodson, E.J., Emsley, P., Evans, P.R., Keegan, R.M., Krissinel, E.B., Leslie, A.G.W., McCoy, A., McNicholas, S.J., Murshudov, G.N., S. Pannu, N., Potterton, E.A., Powell, H.R., Read, R.J., Vagin, A., and Wilson, K.S. (2011). Overview of the CCP4 suite and current developments. Acta Crystallogr. D 67, 235–242.

Yang, R., Wong, Y.H., Nguyen, G.K.T., Tam, J.P., Lescar, J., and Wu, B. (2017). Engineering a catalytically efficient recombinant protein ligase. J. Am. Chem. Soc. 139, 5351–5358.

Yu, Y., and Lutz, S. (2011). Circular permutation: a different way to engineer enzyme structure and function. Trends Biotechnol. 29, 18–25.

Zand, R., Agrawal, B.B., and Goldstein, I.J. (1971). pH-dependent conformational changes of concanavalin A. Proc. Natl. Acad. Sci. USA 68, 2173–2176.

Zauner, F.B., Elsässer, B., Dall, E., Cabrele, C., and Brandstetter, H. (2018a). Structural analyses of *Arabidopsis thaliana* legumain gamma reveal the differential recognition and processing of proteolysis and ligation substrates. J. Biol. Chem. 293, 8934–8946.

Zauner, F.B., Dall, E., Regl, C., Grassi, L., Huber, C.G., Cabrele, C., and Brandstetter, H. (2018b). Crystal structure of plant legumain reveals a unique two-chain state with pH-dependent activity regulation. Plant Cell 30, 686–699.

Zheng, H., Cooper, D.R., Porebski, P.J., Shabalin, I.G., Handing, K.B., and Minor, W. (2017). CheckMyMetal: a macromolecular metal-binding validation tool. Acta Crystallogr. D Struct. Biol. 73, 223–233.

